# Geothermal ecosystems on Mt. Erebus, Antarctica, support diverse and taxonomically novel biota

**DOI:** 10.1101/2024.06.06.597832

**Authors:** Trine Bertram Rasmussen, Stephen E. Noell, Craig W. Herbold, Ian A. Dickie, Roanna Richards-Babbage, Matthew B. Stott, S. Craig Cary, Ian R. McDonald

**Author notes:** Address correspondence to Ian R. McDonald: Te Aka Mātuatua – School of Science, Te Whare Wānanga o Waikato – University of Waikato, Hamilton, New Zealand.

## Abstract

Mt. Erebus, Antarctica, is the southernmost active volcano in the world and harbors a diverse and geochemically unique array of geothermal ecosystems including ‘Subglacial’ and ‘Exposed’ features, surrounded by a vast desert of ice and snow. Previous studies, although limited in scope, have highlighted the unique and potentially endemic biota present on Mt. Erebus. In this study, we provide a systematic biodiversity study across all domains of life and all types of geothermal features present on Mt. Erebus. We present physicochemical and biological data from 39 Exposed samples and 9 Subglacial samples from Mt. Erebus. The taxonomic novelty of prokaryotes and fungi found supports past hypotheses of high endemism among the biota of Mt. Erebus; in particular, the large number of taxonomically divergent fungal sequences was surprising. We found that different site types had unique physicochemistry and biota; in particular, Exposed sites were significantly warmer than Subglacial sites (median: 40 vs 10℃ for Exposed and Subglacial, respectively) and tended to have greater abundances of photosynthetic organisms (*Cyanobacteria* and *Chlorophyta*). Subglacial sites were characterized by a greater abundance of prokaryotes from the phylum *Actinobacteriota*, correlated with the greater concentrations of Ca, Mg, and Sr present. Additionally, we found that Tramway Ridge differed from other Exposed sites as well as all Subglacial sites in physicochemistry (significantly greater conductivity, water content, total carbon, and total nitrogen levels) and biota (greater relative abundances of order *Nitrososphaeria* and phylum *Bacteroidota*). In this study, we provide a blueprint for future work aimed at better understanding the novel biota of Mt. Erebus.

## Introduction

Mt. Erebus, located on Ross Island, Antarctica, stands as the southernmost active volcano on Earth and the sole known active volcano harboring a permanent phonolite lava lake (1–3). The sustained volcanic activity of Mt. Erebus gives rise to diverse geothermal features, including ice caves, fumaroles, and exposed warm soils, supporting a surprisingly rich diversity of life (4–8). These habitats possess unique characteristics. First, they are profoundly influenced by local volcanic activity, with CO_2_-rich fumarolic gases traversing highly altered soils, depositing minerals and freely available water, thereby altering soil pH (4,9–11). Second, situated in Antarctica, the coldest, driest, and windiest continent on Earth, these isolated geothermal hot soils serve as oases of heat, reductant, and liquid water amid a vastly different, extremely cold, and dry environment. The closest similar geothermal sites, Mt. Melbourne and Mt. Rittman, are 359 and 456 km away to the North respectively (12).

The interplay of hot soils and steam with cold air results in a variety of unique geothermal features such as ice towers, ice hummocks, ice caves, and bare hot soils. Generally, the geothermal areas are categorized as ‘Exposed’ (rarely covered by ice) or ‘Subglacial’ (covered by a ceiling of ice for most of the year). Many pockets of ice-free soil can be clustered into “Hot” sites, such as Western Crater and Tramway Ridge, the latter of which is designated an Antarctic Specially Protected Area (**Fig 1**). The unique nature and diversity of geothermal soils and associated features on Mt. Erebus, combined with its isolation, are predicted to result in a wide range of novel and potentially endemic biota (5) and are also predicted to serve as refugia for life during glacial cycles (13). Previous biodiversity studies on Mt. Erebus have been constrained in scope, often having a narrow sampling scale and/or relying on culture (for prokaryotes/fungi) or direct enumeration-based (for non-fungal eukaryotes) methods (10,14–18), both of which are known to underestimate the true diversity present across all domains of life (19).

**Fig 1.**
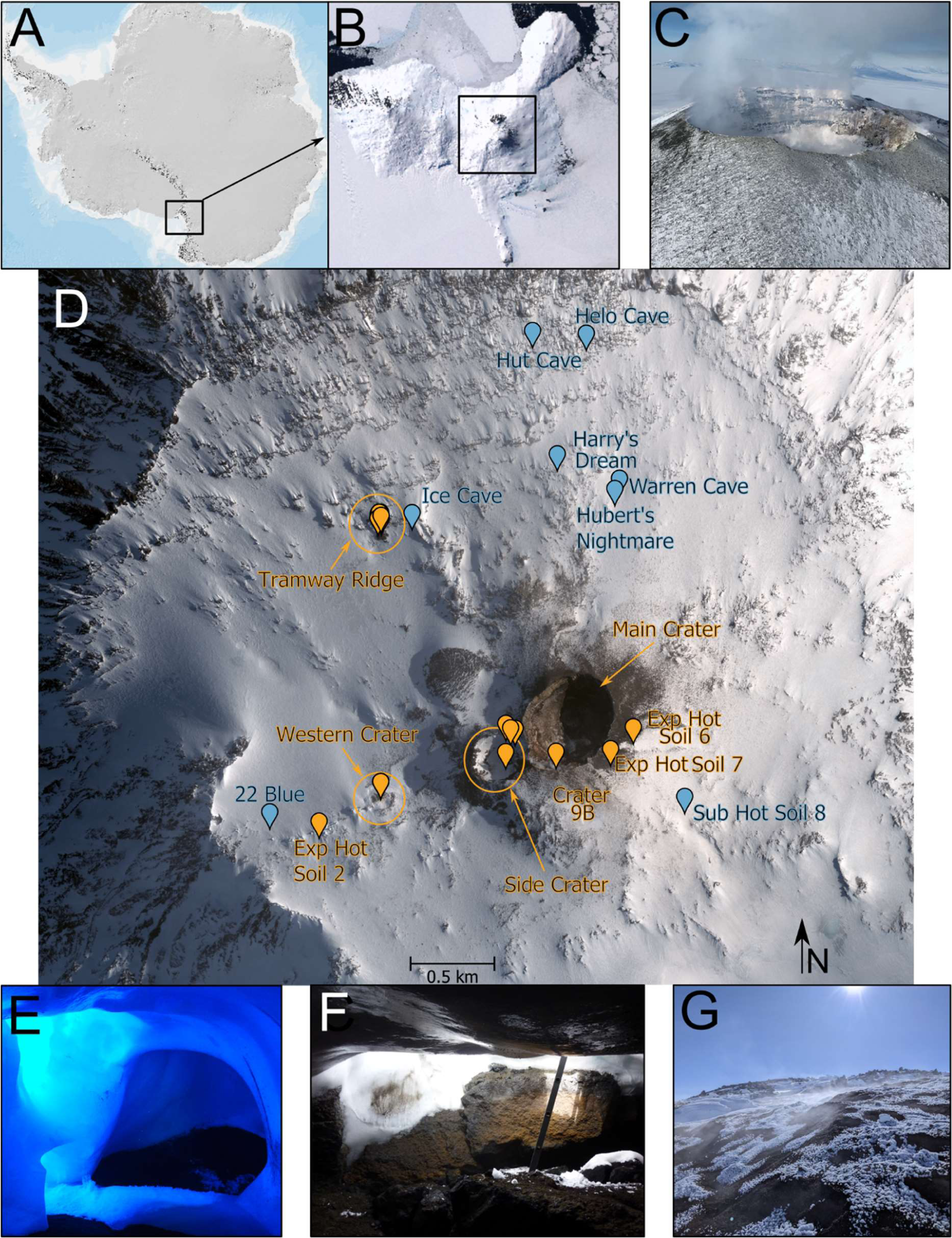
Overview of sampling sites on Mt. Erebus. (A) Map of Antarctica; area enlarged in (B) is shown in the black box. (B) Ross Island, Victoria Land, Antarctica. The location of Mt. Erebus is denoted in the black box. (A-B) are from the Antarctic Digital Database Map Viewer https://www.add.scar.org/, Open Source. (C) Main Crater of Mt. Erebus, harboring a permanent lava lake. (D) Satellite image of Mt. Erebus, showing sampling locations. Sites where more than two samples were collected are indicated with circles. Satellite image was purchased from DigitalGlobe Incorporated, Longmont, CO, USA (2019). (E-G) Representative images of selected sampling sites. (E) Entrance to 22 Blue Cave, as well as the sampling site (mostly dark). (F) Entrance to Helo Cave. (G) Geothermally heated soil at Tramway Ridge. Images in are courtesy of Ian McDonald, Craig Cary, Stephen Noell, and Matthew Stott.

For prokaryotes, culture-independent (genetic-based) studies on Mt. Erebus have been limited to only two Exposed hot soil sites (Tramway Ridge and Western Crater) and three Subglacial sites (Hubert’s Nightmare, Harry’s Dream, and Warren Cave). At the Exposed Tramway Ridge and Western Crater sites, contrasting communities of primarily chemoautotrophic prokaryotes driven by differences in soil pH were described (7). In particular, past studies of Tramway Ridge features have revealed mainly cosmopolitan thermophilic prokaryotes, including many culturable heterotrophs, at the surface, with increasingly divergent lineages found a few centimeters into the soil (4,5,10,14,15,20). At Subglacial sites, a single past study found that autotrophic CO and CO_2_ fixation was a major functional trait of the prokaryotes present (8,21). For fungi, culture-independent studies are limited to Subglacial sites, revealing a diversity of fungal strains from *Basidiomycota* and *Ascomycota* (6,22), including a high dominance of *Malassezia*, which is often human-associated (6) but is also widespread in environmental samples (23,24) and a known contaminant in PCR reagents (25). Culture-based fungal studies have resulted in only a few thermotolerant (26) fungi being isolated from Tramway Ridge (10,27). For non-fungal eukaryotes, only one culture-independent study has been conducted on Mt. Erebus, finding evidence for the presence of arthropods, oligochaetes, and nematodes at a few Exposed and Subglacial sites (6). Past observational studies have noted mosses and epiphytic green algae, with reports of soil fauna confined to limited sightings of amoebae and rotifers inhabiting the moss beds at Tramway Ridge below 40 °C (16,17,28). Despite this past work on Mt. Erebus, there remains no single survey using consistently applied culture-independent methodologies across all three domains of life.

In this study, we conducted an in-depth survey of the Mt. Erebus biota across the three domains of life, encompassing almost all described geothermal sites with the goal of better understanding the microbial ecology of this unique ecosystem. To address this goal we proposed three hypotheses: (1) that the two different site categories, Subglacial and Exposed, would have significantly different biological communities (in terms of community composition and potential functions) that correlate to physicochemical factors, and that Tramway Ridge would differ from other Exposed samples in physicochemistry and biological community, based on past results (7); (2) that we would find a large number of taxonomically novel species, a potential marker of endemism; (3) and that, recognizing the potential influence of human activity on these sites, Subglacial samples would have the highest signature of potential human impact (22). To test these hypotheses, we employed 16S rRNA, 18S rRNA, and ITS gene sequencing, alongside a diverse range of physicochemical measurements. This study represents the first comprehensive, culture-independent survey of life at geothermal features on Mt. Erebus.

## Results

### The physicochemistry of Mt Erebus Subglacial and Exposed soils differ

Despite all samples sites being within 3 km of each other, the physicochemical conditions of geothermal features across Mt. Erebus varied greatly, with a Principal Coordinates Analysis (PcoA) showing that Subglacial samples tended to cluster separately from Tramway Ridge samples, which were more scattered (**S3 Fig**). Other Exposed samples (from here, ‘Exp. Hot Soils’) either clustered more with Tramway Ridge samples or Subglacial samples, with the latter being characterized by elevated Ca, Mg and Sr concentrations, similar to those found in the Subglacial samples (**S3 Fig**). This prompted us to further analyze the physicochemical data with Tramway Ridge samples split from Exp. Hot Soils samples.

Many physicochemical parameters showed significant (Dunn’s test, adjusted *p*-value < 0.05, Benjamini-Hochberg correction) differences between the three site subcategories (**Fig 2**). Exp. Hot Soils and Tramway Ridge samples were distinguishable from Subglacial samples by having higher temperatures (medians: 51, 31, and 7.4 ℃ for Tramway Ridge, Exp. Hot Soils, and Subglacial, respectively) (**Fig 2**). Most other parameters (conductivity, total carbon, total nitrogen, water content, and Cd) were significantly higher at Tramway Ridge compared to all other site subcategories (**Fig 2**). Notably, Tramway Ridge had significantly lower median values for Mg (2406, 6299, 5514 ppb for Tramway Ridge, Exp. Hot Soils, and Subglacial, respectively), Ca (4894, 9269, and 12358 ppb for Tramway Ridge, Exp. Hot Soils, and Subglacial, respectively), and Sr (54.2, 104.2, and 192.0 ppb for Tramway Ridge, Exp. Hot Soils, and Subglacial, respectively) compared to other site subcategories. Tramway Ridge samples also tended to be more acidic than other samples, although this difference was only significant for Exp. Hot Soils compared to Tramway Ridge (medians: 4.75 and 6.93 for Tramway Ridge and Exp. Hot Soils, respectively). Because of the clustering of Tramway Ridge samples from other Exposed samples in the PCoA and the clearly distinct physicochemistry of Tramway Ridge from other Exposed samples, we chose to divide our “Exposed” category into “Tramway Ridge” and “Exposed Hot Soil” subcategories for the biological analyses.

**Fig 2.**
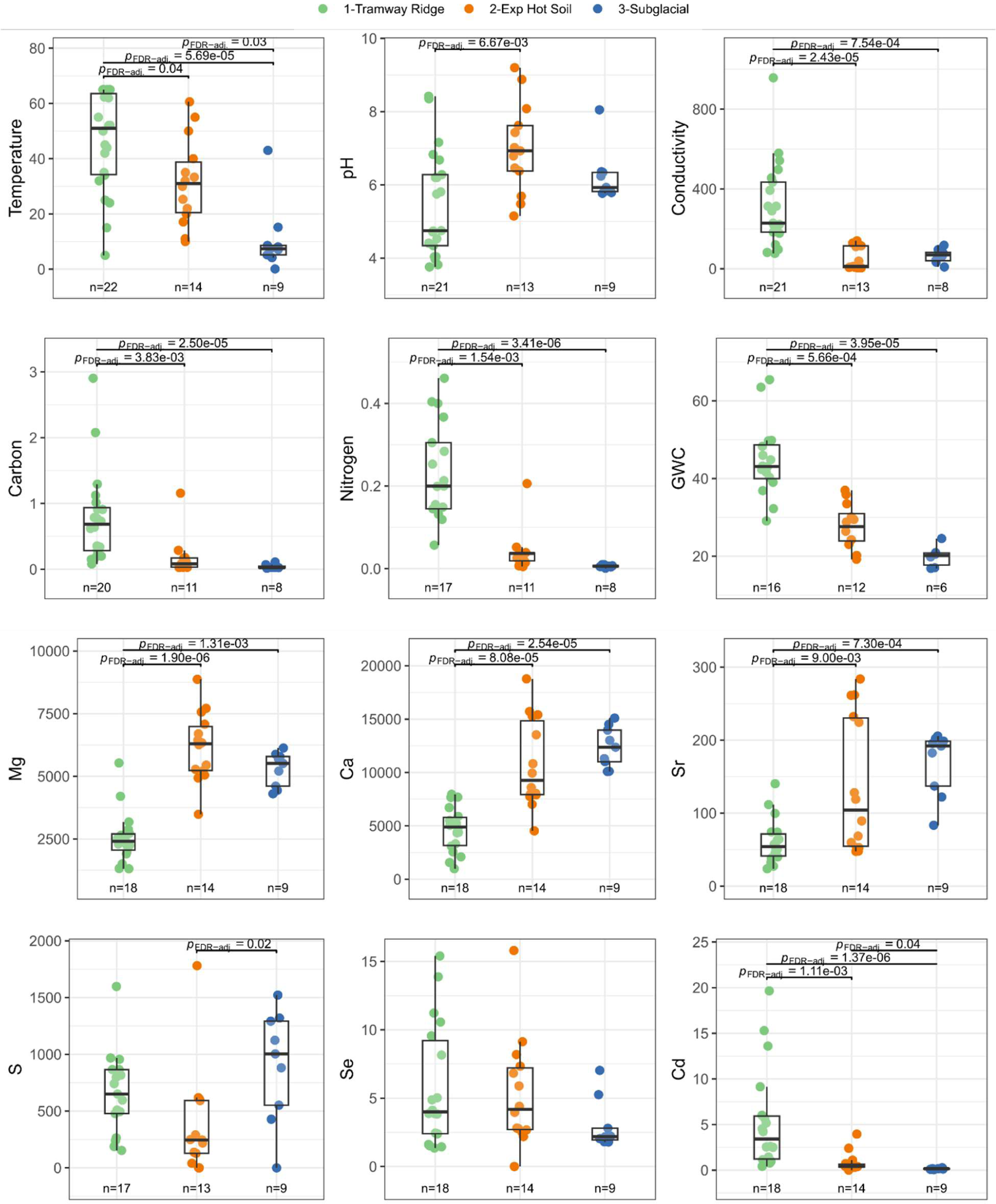
The range of select measured physicochemical parameters within Tramway Ridge (green), Exposed Hot Soils (orange), or Subglacial (blue) from Mt. Erebus geothermal soils. Only parameters that were significantly (two-sided t-test, Benjamini-Hochberg adjusted *p*-value < 0.05) different between Subglacial and all Exposed samples. *p*-values given in the figure (adjusted via a Benjamini-Hochberg correction) are from a Dunn’s Test, following a significant result from a Krustal-Wallis test. Only significant *p*-values are shown. GWC = gravitational water content. Number of tests performed: 12. Data presented has outliers removed, as specified in the Methods.

### Initial analysis of biological communities reveals large proportions of unclassified microorganisms

To assess the diversity of microorganisms inhabiting these sites, we determined the microbial community composition via amplicon sequencing of the 16S rRNA gene (prokaryotes), ITS gene (fungi), and 18S rRNA gene (non-fungal eukaryotes). We observed 2,734 prokaryotic ASVs, 214 fungal ASVs, and 512 non-fungal eukaryote ASVs (see **Tables S2, S3, and S4** for ASV taxonomies and abundance profiles). Of note, a substantial proportion of ASVs in all three communities appeared to be novel with 59.4 % and 86% of prokaryotic and non-fungal eukaryotic taxa unable to be assigned to genera respectively (5.3 % and 6 % of ASVs could not be assigned to a prokaryotic and non-fungal eukaryotic phylum).

For the fungal community, we found many ASVs that were taxonomically unclassified and poorly resolved phylogenetically, which led to a refinement of phylogenetic methodology. This analysis identified 87 clades (>50% support on phylogenetic tree branch) of fungi within a large polytomy (clades with <50% support were defined as polytomies; it was expected that the basal phylogeny would not be resolved given the limitations of the ITS gene region; **S4 Fig**). 46 clades were comprised of a single ASV, 17 clades had two ASVs, and the remainder contained up to 14 ASVs. Based on BLASTn sequences that overlapped 75% or more of the query length, 45 clades could be tentatively identified based on >90% similarity to a known reference sequence in the UNITE database and six additional clades matched a sequence in the wider UNITE+INSD database. An additional 10 clades matched at least one sequence in the NCBI database, with six of these only matching a single sequence at > 90% identity. In total, 20 clades had no matching sequences in any database at > 80% identity (**Table S5**). While most of these clades had only a single ASV, one contained 5 ASVs and another contained 3.

### Differential abundances of several taxonomic groups distinguish site types

Unlike in the physicochemical results, we found that mean community alpha diversity (measured via the Shannon index) was similar across site categories, although there was substantial variation within site categories for the prokaryotes and non-fungal eukaryotes (**Fig 3A, C, E**). Mean community Shannon diversity tended to be highest across prokaryotes (3.5-3.75), then non-fungal eukaryotes (1.6-2.2), then fungi (1.7).

**Fig 3.**
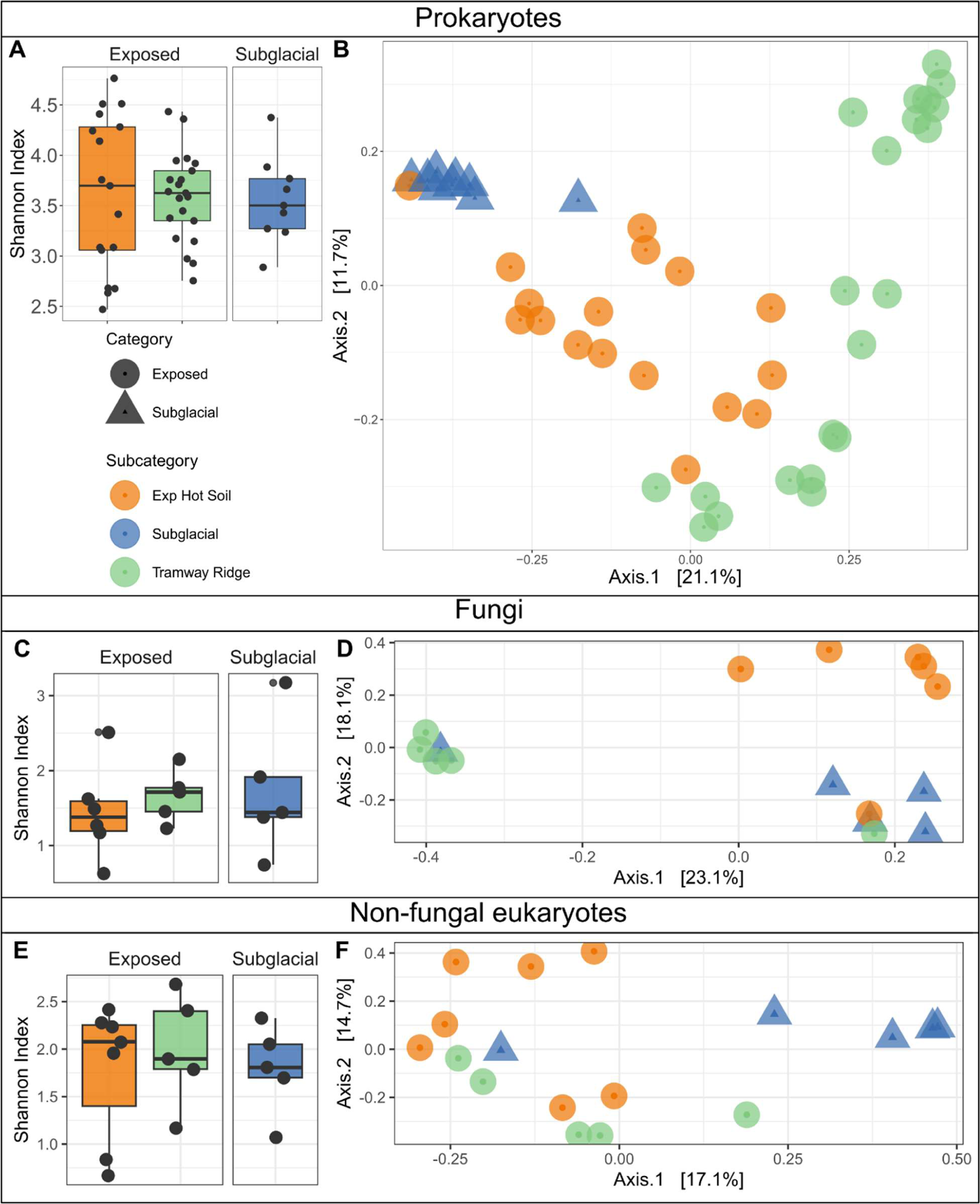
Biodiversity analysis of prokaryotic (16S rRNA gene), fungal (ITS) and non-fungal Eukaryote (18S rRNA gene) datasets across Mt Erebus Exposed and Subglacial geothermal sites. (A, C, E) Alpha diversity measured with the Shannon index. (B, D, E) Beta diversity measured with a PCoA using Unifrac distances.

However, in our beta diversity analysis (**Fig 3**) using PCoA, we found that, among prokaryotic, fungal, and non-fungal eukaryotic data sets, Subglacial and Exposed samples generally clustered separately from each other, although to different extents depending on the data set (**Fig 3B, D, F**). This trend was confirmed with significant PERMANOVA test results (*p* = 0.001) and non-significant results for different dispersions between subcategories for both the fungal and non-fungal eukaryotic data sets. Exposed hot soil samples were significantly different both from Tramway Ridge samples (*p* = 0.009, 0.024 for fungal and non-fungal eukaryotes, respectively) and from Subglacial samples (*p* = 0.01, 0.016 for fungal and non-fungal eukaryotes, respectively), but Tramway Ridge and Subglacial samples were not significantly different (*p* = 0.12, 0.06 for fungal and non-fungal eukaryotes, respectively). Due to the significantly different dispersions of the samples, a reliable PERMANOVA test for significant differences in prokaryotic communities between sample categories or subcategories was not feasible. In the prokaryotic data set, it was especially notable that two distinct clusters emerged within Tramway Ridge samples, with high-temperature samples (>55°C) tending to cluster separately. Also of note was the clustering of one Exposed hot soil sample (Crater 6 low) with the Subglacial samples in both prokaryotic and physicochemical data.

To identify the taxonomic groups responsible for the differential clustering of the site categories, we used a differential abundance test within the microeco package in R and an indicator species analysis. In the prokaryotic data, our differential abundance test identified several phyla that had significantly higher relative abundance averages in Exposed samples: *Crenarchaeota* (especially class *Nitrososphaeria*), *Deinococcota*, *Armatimonadota*, *Cyanobacteria*, *Nitrospirota, Firmicutes,* and GAL15 (adjusted *p* < 0.05, Benjamini-Hochberg correction) (**Fig 4A, B, 5A**). Conversely, the average relative abundances of *Actinobacteriota* (mainly classes *Actinobacteria* and *Thermoleophilia*)*, Chloroflexi* (especially classes AD3 – now candidate phylum *Ca.* Dormibacteriota – and *Ktedonobacteria*)*, Acidobacteria*, and *Alphaproteobacteria* were significantly greater in Subglacial sites (adjusted *p* < 0.05, Benjamini-Hochberg correction) (**Fig 5A, S4A**). Exposed hot soils differed from Tramway Ridge by having significantly higher relative abundances of *Firmicutes* and *Actinobacteriota*, although the latter had a high variance in relative abundance at Exposed hot soils (from <1 % up to 32.9 % in Crater 8 < 3cm) (**Fig 5B**). Our indicator species analysis at the ASV level confirmed these broad trends, with a few nuances (**Table S6**). Of particular note was the finding that the genus *Fischerella* is an indicator for Tramway Ridge, while *Leptolyngbya* is an indicator more broadly for Exposed sites; also, we also found four indicator species for Tramway Ridge that were from the phylum *Bacteroidota*, which was not identified in the differential abundance analysis (**Table S6**).

**Fig 4.**
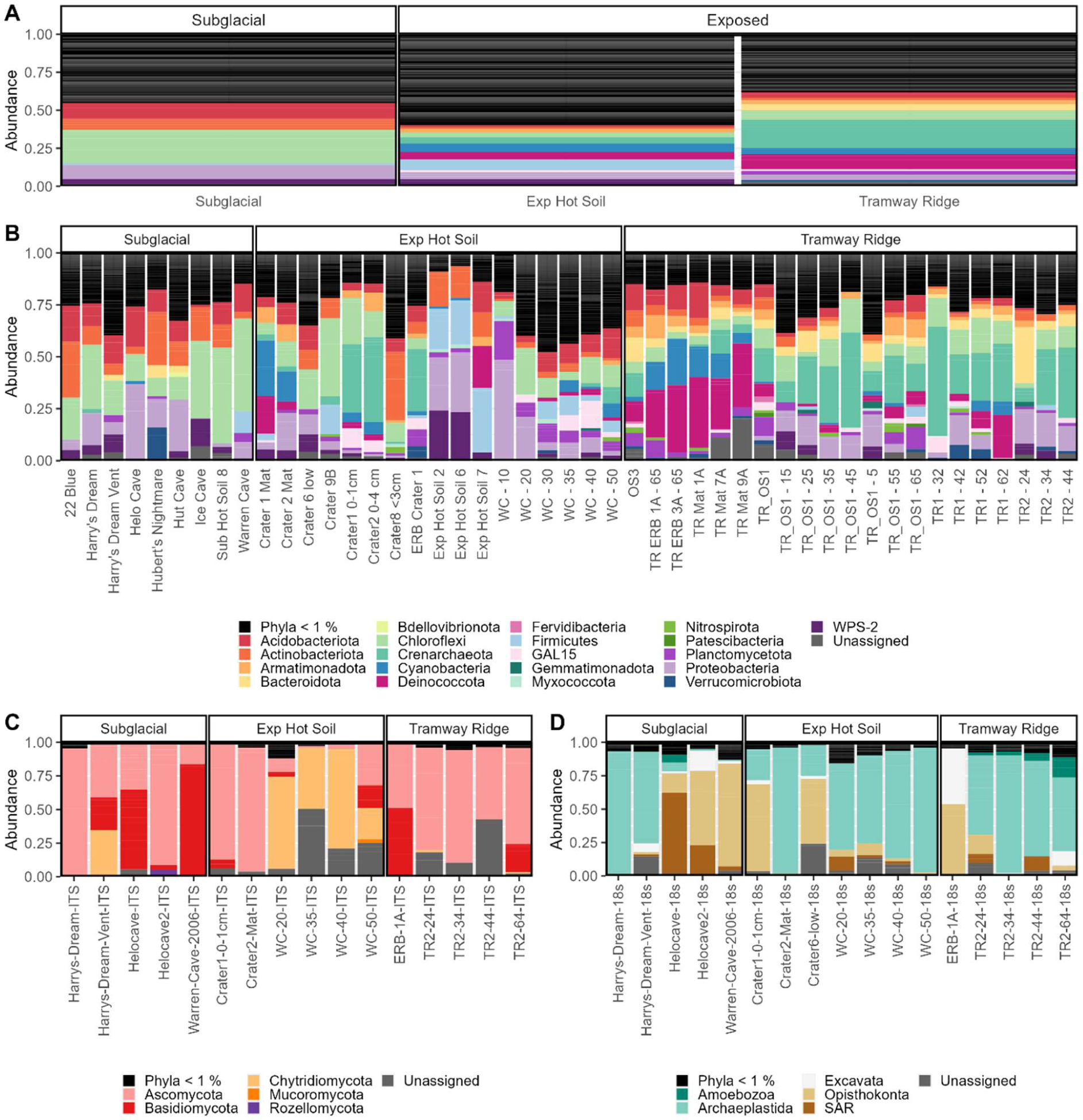
Microbial community composition shown as relative abundances of prokaryotic (A-B) and eukaryotic phyla (C-D). The average relative abundance of prokaryotic phyla in subglacial, exposed hot soils, and Tramway Ridge samples are seen in A, while B shows the relative abundances for each sample individually. Similarly, C and D shows the relative abundances of fungal and other eukaryotic phyla in individual samples.

**Fig 5.**
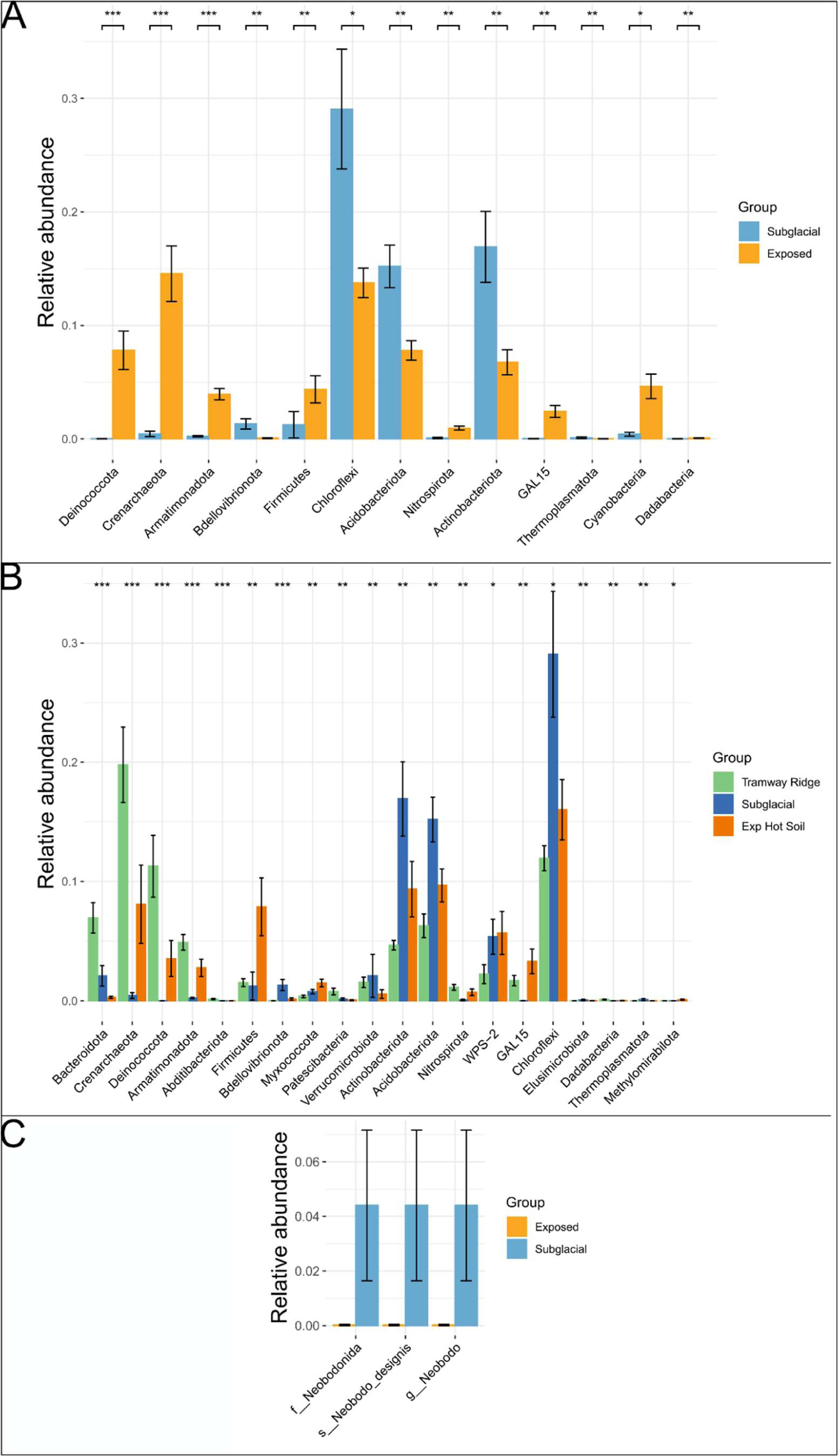
Results of a differential abundance test. (A-B) Prokaryotic phyla that significantly differed in relative abundances between (A) Subglacial vs. Exposed samples or (B) all three subcategories. (C) Taxa at any rank that were significantly different between Exposed and Subglacial sites in the non-fungal eukaryotic data set. For all, significant difference in abundance was determined using the random forest method within the microeco package. The significance stars correspond to the following Benjamini-Hochberg adjusted *p*-values: * = 0.05, ** = 0.01, *** = 0.001, **** = 0.0001.

For the fungi, at the phylum level, we found that the three site subcategories were generally distinguishable by the dominance of phyla (**Fig 4C**). For Subglacial samples, ASVs from *Ascomycota* and *Basidiomycota* were predominant in most samples. Samples from Western Crater were distinguishable by very high abundances of ASVs from *Chytridiomycota*, while Tramway Ridge samples were almost completely dominated by ASVs from *Ascomycota*. At taxonomic levels below phylum, conclusions were challenging to draw due to the large number of ASVs that could not be assigned even to a class (**S5 Fig B**). However, when examining the distribution patterns of different clades (generated from our phylogenetic approach), we found that the most highly divergent clades (that could not be reliably assigned at phylum or class level; e.g., clades 1, 3, 4, 5, and 42) were found in Exposed samples (e.g., clade 1, predominantly found at Western Crater, is a highly divergent member of *Chytridiomycota*), while clades that were most abundant in Subglacial samples were from better characterized groups (e.g., clade 32 – *Leotiomycetes*, clade 14 – *Agaricomycetes*, etc.) (**S6 Fig**). In our differential abundance analysis, there were no fungal taxonomic groups identified as significantly different between sample categories. However, our indicator species analysis identified four ASVs from *Ascomycota* as indicators for Tramway Ridge, two of which were from the Family *Herpotrichiellaceae* (**Table S6**).

At a broad level, the non-fungal eukaryotic data was dominated by ASVs from the *Archaeplastida* group (*Chloroplastida*), apart from three Subglacial samples (two Helo cave samples, one Warren Cave sample). These three Subglacial samples were almost completely lacking in *Archaeplastida* ASVs, instead being dominated by ASVs from the SAR group and non-fungal *Opisthokonta*. The *Chloroplastida* ASVs detected were mainly from green algal groups (*Pseudochlorella, Coccomyxa, Coenocystis,* etc.) or moss chloroplasts. Non-photosynthetic eukaryotes detected included alveolates (mainly ciliates), amoeboids (e.g., the Cercazon *Gymnophrys*, and Amoebozoan *Tubulinea, Vermamoeba,* and *Flamella*), and other protists (e.g., the Excavata *Neobodo*). In our differential abundance analysis, only one taxonomic group was identified as significantly different between the three site categories: the species *Neobodo designis*, which was present only at Subglacial sites at 4% relative abundance (**Fig 5C**). Again, however, our indicator species proved more sensitive, identifying a number of interesting indicator species; of note were four *Chloroplastida* ASVs (two each at Tramway Ridge and Exposed Hot Soils), ASVs from the genera *Monodus* and *Didinium* in Exposed Hot Soils samples (yellow-green unicellular algal genus and carnivorous ciliate, respectively), *Neobodo designis* at Subglacial samples, and ASVs from the genera *Tubulinea* and *Gymnophrys* (amoeba and predatory amoeboid Cercazoan, respectively) (**Table S6**).

### Network analysis

We used a co-occurrence network to investigate whether co-occurrence patterns across all domains of life also point towards partitioning of Erebus biota into the three site subcategories, Subglacial, Tramway Ridge, and Exposed Hot Soils. Based on our other results, we expected that ASVs from each of the three site subcategories would show greater interconnectedness within subcategory than to other subcategories, resulting in clustering of ASVs based on site subcategory. Abundances of ASVs within each of the three datasets were center-log-ratio transformed (CLR) prior to joining data sets to attempt to ameliorate compositional bias, and ASVs were then trimmed by abundance to 228 ASVs (>4.5 CLR in at least three samples). The resulting co-occurrence network was divided into four clusters by the fast greedy modularity optimization algorithm, one of which only contained one ASV (**Fig 6A**). The modularity of the network was 0.43, indicating medium connectivity; 85.6% of the connections indicate positive correlations.

**Fig 6.**
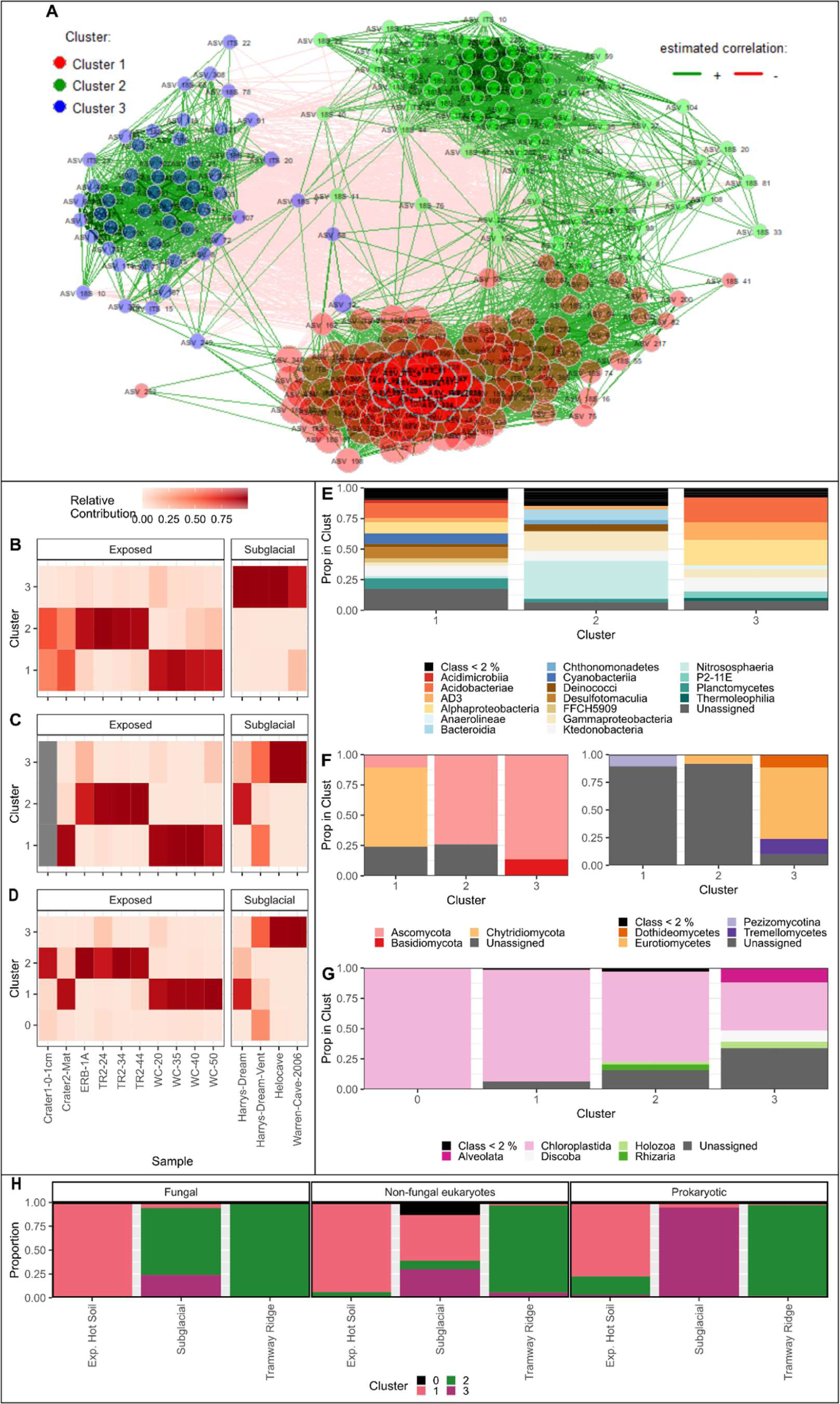
Co-occurrence network to investigate abundance patterns in the abundant eukaryotes, non-fungal eukaryotes and prokaryotes. (A) Co-occurrence network using Pearson correlations, where each dot is an ASV, colored by cluster; ASVs within clusters are more connected to each other than to ASVs outside of the cluster. (B - D) Heatmaps showing the relative abundance of each cluster at each site for (B) prokaryotes, (C) fungi, and (D) non-fungal eukaryotes. (E - G) Bar graphs showing what proportion of each cluster belongs to what class for (E) prokaryotes, (F) fungi, or (G) non-fungal eukaryotes. For (F), proportion of cluster belonging to fungal phyla is shown on the left and class on the right, given the large proportion of fungal ASVs that were unassigned at the class level. (H) Bar plot showing the contribution of ASVs from each cluster to the total number of reads in each site subcategory for each type of organism.

Based on the total abundance of reads from each cluster, we found support for our hypothesis of partitioning based on site subcategory, with each of the clusters generally corresponding to one of three site subcategories: Exp. Hot Soils (Cluster 1), Tramway Ridge (Cluster 2), or Subglacial (Cluster 3) (**Fig 6B-D, H**). In general, ASVs from one of the three clusters comprised over 80% of the reads in any given subcategory. However, this was not the case for the fungal and the non-fungal eukaryotes, where ASVs from clusters other than Cluster 3 had strong contributions to Subglacial communities (**Fig 6C-D, H**). However, this was primarily only at Harry’s Dream, which is known to be a cave with greater light levels (8).

We also observed that Clusters 1 and 2 were more connected to each other than to Cluster 3 (162 positive connections between 1 and 2, vs. 39 and 8 for Clusters 3 and 1 and Clusters 3 and 2, respectively), indicating more similarity in co-occurrence patterns within Exposed sites than between Exposed and Subglacial sites (**Fig 6A**). The composition of the clusters generally matched our community composition analyses (**Fig. 4**); in particular, Cluster 3 had few photosynthetic organisms (*Cyanobacteria* or *Chloroplastida* in the 16S or 18S rRNA gene surveys, respectively), Clusters 1 and 2 had much larger proportions of novel fungi at the class level than Cluster 3, and Cluster 2 prokaryotes were enriched in *Nitrososphaeria* and *Deinococcus* (**Fig 6E-G**).

### Do environmental factors correlate with microbial community composition?

We next looked at what physicochemical factors might be influencing the different microbial communities observed at the three site types using a Mantel test and a distance-based redundancy analysis (db-RDA). However, we first observed that several factors were strongly correlated with each other across all site types, especially Mg, Ca, and Sr, as well as total carbon and nitrogen (**Table S7**). As autocorrelation can strongly impact the results of Mantel tests, we chose to remove Sr from this analysis as it is not known to have any strong relationship with biotic communities. We then removed either Mg or Ca, or total carbon or nitrogen, and re-ran the Mantel test, with the finding that the correlation coefficients and adjusted *p*-values for all factors remained almost identical between all tests, indicating that our findings are robust to autocorrelation (**Table 1**).

**Table 1.** Physicochemical factors that correlate with prokaryotic communities determined via a Mantel test (Spearman correlation on Bray-Curtis distance matrix between biotic communities) to identify. Only factors with significant adjusted *p*-values are shown. Test was re-run twice with autocorrelated factors Sr and one of Mg or Ca removed. Adjusted *p*-values are the result of a Benjamini-Hochberg adjustment for multiple comparisons; number of tests performed: 27 prior to removing any factors, 25 after removing Sr and Ca or Mg, 26 after removing Total Carbon or Nitrogen. GWC: gravitational water content.

| Variables | Correlation coefficient | *p*-value | Adjusted *p*-value |
| --- | --- | --- | --- |
| Temperature | 0.350658773 | 0.002 | 0.0108 |
| pH | 0.198262798 | 0.004 | 0.0135 |
| Conductivity | 0.208247108 | 0.003 | 0.011571429 |
| Carbon | 0.212146947 | 0.005 | 0.015 |
| Nitrogen | 0.273662297 | 0.003 | 0.011571429 |
| GWC | 0.332289989 | 0.001 | 0.00675 |
| Mg | 0.411655232 | 0.001 | 0.00675 |
| Ca | 0.464576911 | 0.001 | 0.00675 |
| Sr | 0.41058035 | 0.001 | 0.00675 |
| **Re-run without Sr and Mg** |  |  |  |
| Temperature | 0.350658773 | 0.001 | 0.008333333 |
| pH | 0.198262798 | 0.006 | 0.025 |
| Conductivity | 0.208247108 | 0.005 | 0.025 |
| Carbon | 0.212146947 | 0.008 | 0.028571429 |
| Nitrogen | 0.273662297 | 0.002 | 0.0125 |
| GWC | 0.332289989 | 0.001 | 0.008333333 |
| Ca | 0.464576911 | 0.001 | 0.008333333 |
| **Re-run without Sr and Ca** |  |  |  |
| Temperature | 0.350658773 | 0.001 | 0.008333333 |
| pH | 0.198262798 | 0.008 | 0.028571429 |
| Conductivity | 0.208247108 | 0.007 | 0.028571429 |
| Carbon | 0.212146947 | 0.006 | 0.028571429 |
| Nitrogen | 0.273662297 | 0.002 | 0.0125 |
| GWC | 0.332289989 | 0.001 | 0.008333333 |
| Mg | 0.411655232 | 0.001 | 0.008333333 |
| **Re-run without Total Nitrogen** |  |  |  |
| Temperature | 0.350659 | 0.001 | 0.0052 |
| pH | 0.198263 | 0.004 | 0.017333 |
| Conductivity | 0.208247 | 0.005 | 0.018571 |
| Carbon | 0.212147 | 0.008 | 0.026 |
| GWC | 0.33229 | 0.001 | 0.0052 |
| Mg | 0.411655 | 0.001 | 0.0052 |
| Ca | 0.464577 | 0.001 | 0.0052 |
| Sr | 0.41058 | 0.001 | 0.0052 |
| **Re-run without Total Carbon** |  |  |  |
| Temperature | 0.350659 | 0.001 | 0.0052 |
| pH | 0.198263 | 0.005 | 0.018571 |
| Conductivity | 0.208247 | 0.006 | 0.0195 |
| Nitrogen | 0.273662 | 0.002 | 0.008667 |
| GWC | 0.33229 | 0.001 | 0.0052 |
| Mg | 0.411655 | 0.001 | 0.0052 |
| Ca | 0.464577 | 0.001 | 0.0052 |
| Sr | 0.41058 | 0.001 | 0.0052 |

For prokaryotes, analyses revealed that the measured physicochemical factors responsible for the separation of Tramway Ridge from Subglacial and Exposed hot soil samples along the first axis were elevated temperature, nitrogen, carbon, moisture (gravimetric water content, GWC), and conductivity in Tramway Ridge samples (**Fig 7**). Subglacial samples and some Exposed hot soil samples were characterized by elevated concentrations of Ca and Mg. An RDA analysis was conducted to explore the contribution of different phyla in explaining the grouping patterns observed in **Fig 7** (**S7 FigA**). The top ten taxa that explained the most variation in the axes were included as explanatory variables, increasing the percentage of variation explained by the axes (40.1% to 60.7% for the first axis). The phyla *Deinococcota, Bacteroidota,* and *Crenarchaeota,* were found to be associated with samples from Tramway Ridge. Conversely, *Actinobacteriota* were associated with other Exposed hot soils and Subglacial soils. The dispersion of Exposed hot soil and Subglacial soil samples along the second axis appeared to be explained by several phyla: *Proteobacteria*, WPS-2, and *Firmicutes* were associated with Warren Cave, Sub Hot Soil 8, Harry’s Dream, Crater 6 low, Crater 8 < 3cm, 22 Blue, and Western Crater samples. While *Acidobacteria, Planctomycetota,* and *Chloroflexi* were associated with Exp Hot Soil 7, Helo Cave, Hut Cave, Exp Hot Soil 2, and Exp Hot Soil 6. Furthermore, several phyla that contributed to the variation observed in the RDA analysis also exhibited significant correlations with the physicochemical factors (spearman; *p*< 0.05, Benjamini-Hochberg adjustment) (**S7 FigB)**. This provides additional support to the hypothesis that these physicochemical factors play a crucial role in shaping the distinct taxonomic compositions of the different environments.

**Fig 7.**
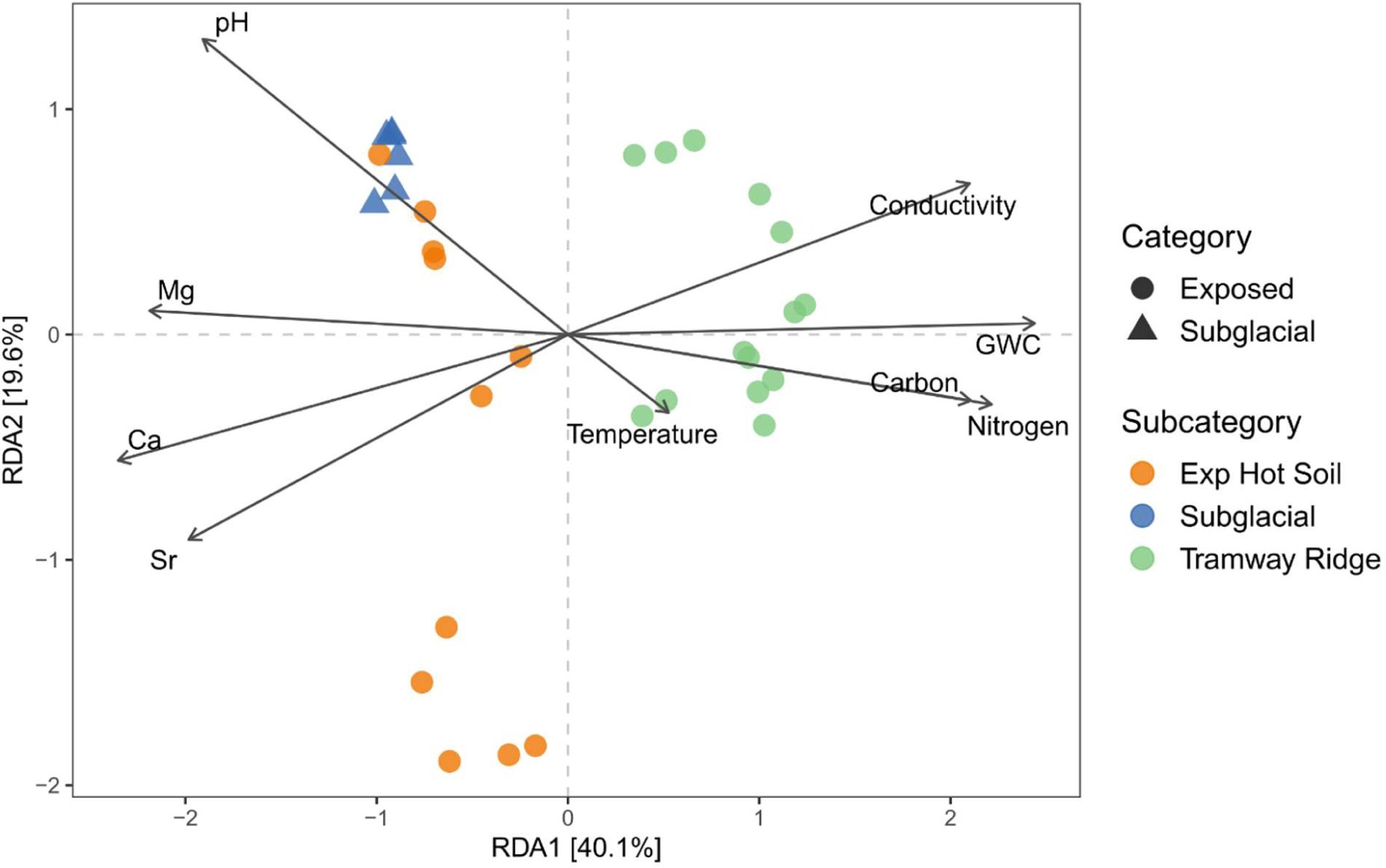
Distance-based redundancy analysis of environmental factors that correlated with the prokaryotic communities. Environmental factors shown had a significant (*p* < 0.05) correlation with the communities based on a Mantel test after adjusting for multiple comparisons using a Benjamini-Hochberg adjustment (see Table 1).

For both the fungal and non-fungal eukaryotic communities, it was more difficult to infer correlations between physicochemical parameters and community compositions due to the small number of samples from each sample category. Indeed, we did not find any physicochemical parameters that significantly (*p* < 0.05, Benjamini-Hochberg correction) correlated with the community compositions using a Mantel test.

### Predicted prokaryotic functional profiles correlate with the physicochemistry of Exposed and Subglacial sites

We used a functional prediction program (FAPROTAX) to investigate correlations between predicted functions and environmental drivers of prokaryote biodiversity, determined above (**Fig 8**). We did not attempt similar analyses for eukaryotic data sets due to the unique fungal taxa detected and the lack of comprehensive functional data sets for non-fungal eukaryotes. Exposed sites differed from Subglacial sites by containing photosynthetic microbes. In contrast, Subglacial sites were predicted to contain nonphotosynthetic *Cyanobacteria*. Communities at Exposed sites were also predicted to contain the potential for nitrification and aerobic ammonia oxidation, while Subglacial sites were predicted to support methylotrophy, methanotrophy, dark sulfide oxidation, chitinolysis, and anaerobic chemoheterotrophy.

**Fig 8.**
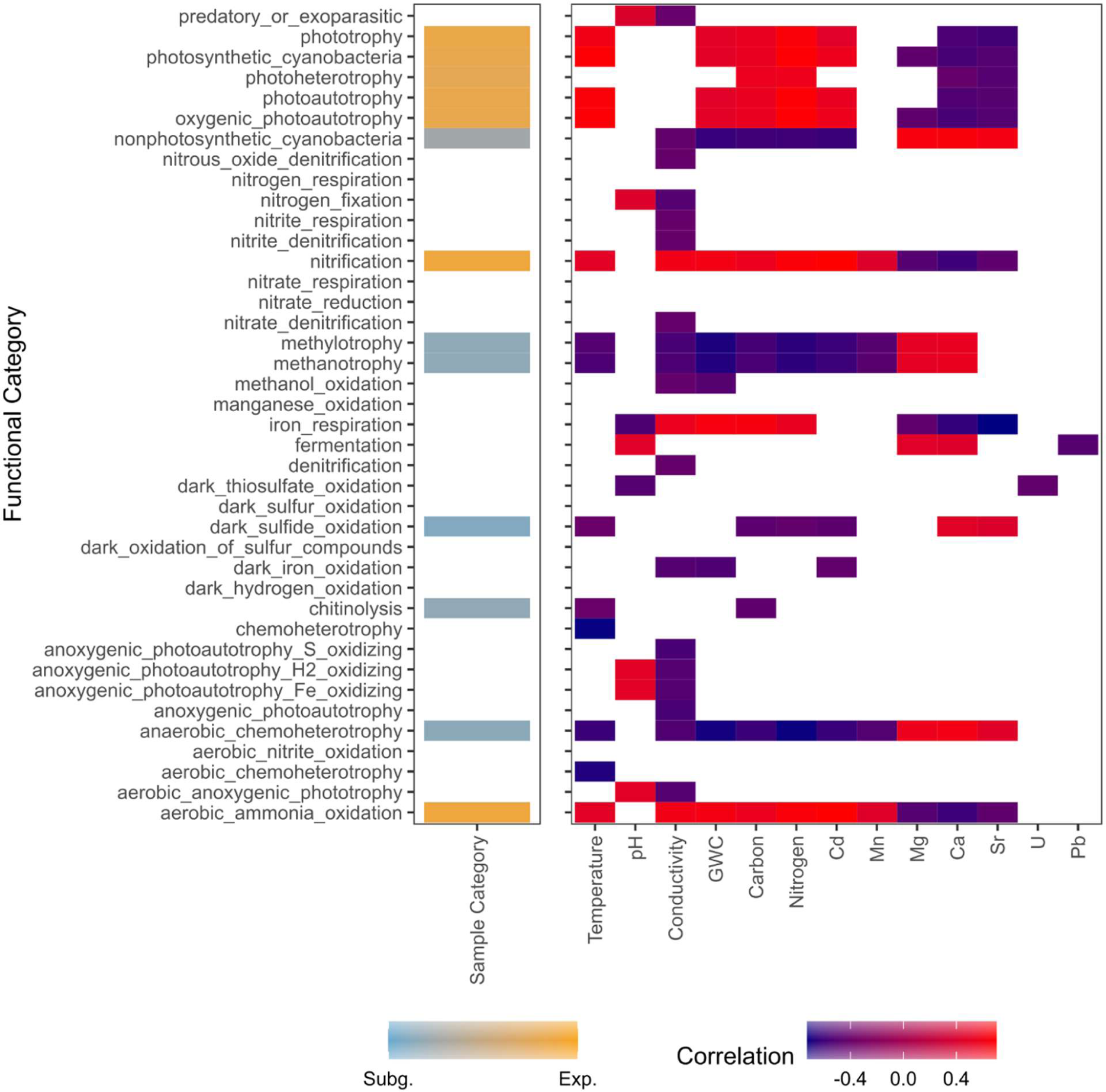
Predicted functional profiles that significantly correlate (spearman’s correlation, Benjamini-Hochberg adjusted *p*<0.05; 784 tests performed) with either Site category (Subglacial vs. Exposed) or physicochemical factors potentially driving different communities across Exposed and Subglacial sites. Non-significant correlations are shown as empty cells.

### Potential Human Impacts on Mt. Erebus Biota

We examined our data set for potential markers of human contamination, specifically to identify if any sites are more impacted than others. Previously, *Malasezzia* yeast species were identified in Warren Cave in 2013 at high abundance (37% relative abundance), with their presence attributed to human contamination (22), although this attribution has been called into question lately (23–25). We did observe *Malasezzia* species in our data set, with lesser abundance (<15% relative abundance) at Helo Cave and one sample each from Western Crater and Tramway Ridge. We found none at Warren Cave, however. We also observed lignolytic fungi in our data set: clade 26 (100% ID, full length match to *Gymnopilus penetrans*) and clade 63 (highly similar match to *Polyporales*, white rot fungi), found at Helo Cave, Western Crater 20°C, and Western Crater 50°C. ASVs from both clades were present only in very low abundance (<2% relative abundance), except for ASV 155, 16% at ERB-1A. We did not find evidence for any known human-related bacteria (e.g., *Staphylococcus*, *Bacteroidales*, or *E. coli*) in our prokaryotic data set. We did observe several 18S ASVs that were identified as human DNA (ASVs 186 and 265). These ASVs were at low abundance in only a few samples (0.8% relative abundance at Helo Cave, 0.1% relative abundance at Western Crater 20°C).

## Discussion

Our investigation of the diversity of biota inhabiting geothermal ice-free soils on Mt. Erebus using culture-independent methods provides the broadest survey to-date of life in Earth’s most remote geothermal site. Our findings provide evidence that the many environments supported by thermal features on Mt. Erebus have provided unique habitats for diverse biotic communities. Additionally, our biological data suggest high levels of taxonomically novel microorganisms inhabiting thermal features on Mt. Erebus. Finally, we show limited evidence of human contamination signatures on Mt. Erebus.

### Hypothesis 1: different habitats on Erebus support distinct biota

We found support for our hypothesis that there are distinct biological communities, functions, and correlated physicochemical parameters at Subglacial, Tramway Ridge, and Exposed sites (**Figs 2-6, 8**). We base this on several pieces of evidence. First, we found that the three Subcategories tended to group together in a PCoA plot of the physicochemical data (**S3 Fig**). Second, we showed that the biological communities are significantly different from each other based on a significant PERMANOVA result (although the results from the prokaryotes are confounded by a significant effect of the data’s dispersion on the outcomes of the test), the general clustering of biological communities based on subcategory in the PCoA plots for all domains (**Fig 3**), and the distinct indicator species for each site subcategory for all domains. Finally, our co-occurrence network found that, across all domains of life, ASVs tended to cluster into groups based on prevalence at one of the three site Subcategories (**Fig 6**). A previous study also found that Tramway Ridge and Western Crater harbored distinct microbial communities (7), but here we extend those results to show that Tramway Ridge has, for the most part, distinct biological communities compared to all other Exposed sites and all Subglacial sites. The exceptions to these trends seem to be mainly driven by unique physicochemistry (as will be discussed below for *Actinobacteriota*) or higher light levels (as will be discussed below for certain Subglacial sites). Below, we discuss the distinguishing physicochemical and biological features of each site subcategory.

#### Distinguishing features of Subglacial sites: Actinobacteriota, elevated Ca/Mg/Sr, and bacteriovores

For our prokaryotic data set, one of our more surprising findings was the strong correlation between large concentrations of calcium (Ca), magnesium (Mg), and strontium (Sr) in Subglacial samples and the presence of *Actinobacteriota* (**S7 Fig**). Ca exhibited the greatest correlation coefficient with the structure of microbial communities (**Table 1**), 2.3 times greater than pH. Indeed, the clustering of Exposed sample Crater-6-low (from Side Crater) with other Subglacial samples, despite its high temperature (60.6 °C), seems to be driven more by mineral content than temperature (**Fig 2**). This sample had high abundances of *Actinobacteriota* (**Fig 3B**) and similar amounts of Mg, Ca, and Sr as Subglacial samples (**S2 Fig**). Because Ca, Mg and Sr are colinear variables, it is impossible to conclude which variable drives the *Actinobacteriota* abundance. However, past research has found that Ca has a significant impact on the growth of many *Actinobacteriota* species (29–32). Ca has been suggested to play an important role in spore formation (29), but most *Actinobacteriota* in our data set are not known to be spore-forming (e.g., genera *Conexibacter*, *Mycobacterium*, and *Crossiella* (33,34)). Ca is also known to be involved in antibiotic resistance in some *Actinobacteriota* (35) and in the production of Ca-dependent antibiotics (36–40). It could be that these antibiotics provide a competitive advantage, but this hypothesis remains to be tested. Alternatively, it might be that the *Actinobacteriota* in Subglacial samples increase the levels of Ca in the soil due to the biomineralization of dissolved Ca ions, a hypothesis that was proposed for some cave *Actinobacteriota*.(41) Interestingly, it has been suggested that this process evolved due to an intracellular toxic effect of elevated Ca ions (42). It is therefore not clear whether high Ca concentrations are beneficial for *Actinobacterial* growth or purely a result of their metabolic activity.

Also from our prokaryotic data set, Subglacial samples were distinguishable by the presence of several distinctive predicted functions. However, predictions of functional profiles based on 16S rRNA sequences should be interpreted with utmost care, given that taxonomy is not always a good predictor of function. Of particular note is the observed presence of nonphotosynthetic *Cyanobacteria* (unclassified families within *Sericytochromatia* or unclassified genera in *Obscuribacteraceae*) enriched in Subglacial samples at low abundances (<1% relative abundance) (**Fig 7**). These are poorly understood lineages that are proposed to be the deepest branching ancestor to photosynthetic *Cyanobacteria* (43), relying on mixed-type fermentation or nitrate metabolism (*Obscuribacteraceae*) (44,45), or aerobic respiration to gain energy (*Sericytochromatia*) (46). They have not previously been reported in geothermal soils or caves (44). We also observed an enrichment in methanotrophy and methylotrophy in Subglacial samples, attributed to alphaproteobacterial ASVs from the genera *Methylocapsa*, *Methylocella,* and *Methylocystis* (**Fig 7**). However, methane concentrations in ice caves on Mt. Erebus are very low, measuring just above atmospheric levels (9). Several possibilities to explain this enrichment of methanotrophs include: a reliance on atmospheric levels of methane (observed for some *Methylocapsa* species (47)), a reliance on H_2_ oxidation (previously discovered in some *Methylocystis* species (48)), or a reliance on other hydrocarbons such as ethane or propane (observed in some *Methylocella* species (49)) whose levels have not been measured, despite reports of natural gas-like smells in some ice caves on Mt. Erebus (9).

For our fungal data set, few meaningful patterns could be observed in the Subglacial samples. However, for the non-fungal eukaryotes, we found that subglacial samples were distinguishable primarily by the presence of *Neobodo designis,* heterotrophic flagellates that are bacteriovores generally found in aquatic environments, but have also been found in soil (50). Their absence from Exposed sites could either indicate a preference for cooler temperatures or that they can find greater concentrations of bacterial prey at Subglacial sites. Similarly, ciliate ASVs (also heterotrophic eukaryotes that generally feed on bacteria) were only present at Subglacial samples (**S5 Fig**), indicating similar constraints on distribution as *Neobodo*.

#### Distinguishing features of Exposed sites: influence of sunlight and higher temperatures

Overall, one of the most apparent differences between Exposed and Subglacial sites is the variation in available sunlight. The influence of this disparity is evident in the presence of photosynthetic *Cyanobacteria*, algae (order *Chlorophyta* and genus *Monodus*), and fungi from the class *Eurotiomycetes*, which is primarily a lichenized class, at only Exposed sites and Harry’s Dream, the cave in our data set with the highest light penetration (8) (**Figs 6, 4A, Table S6**). Many of the prokaryotic cyanobacterial ASVs identified belong to the families *Nostocaceae* and *Leptolyngbyaceae.* These *Cyanobacteria* have been previously found at Tramway Ridge and are renowned for being cosmopolitan and thermophilic (7,44,51,52). The most common algae at Exposed samples in our data set have been observed previously on Mt. Erebus, especially *Chlorella* species (17). Similar to past descriptions of these species, we did not find them in samples with temperatures above 35 °C and found them mainly in Harry’s Dream samples. On the other hand, the *Coccomyxa* ASVs in our data set were primarily found in Exposed samples, especially at the Tramway Ridge 4 °C sample, indicating a higher temperature tolerance than *Chlorella*. For the putatively lichenized fungi, our finding of *Eurotiomycetes* ASVs differs from previous results in Antarctica which primarily identified fungi from *Lecanoromycetes*, another primarily lichenized fungal class (53,54). In our 18S and 16S rRNA gene data sets, we only identified two common photosynthetic partners of lichenized fungi: the green alga *Trebouxia* and the cyanobacterium *Nostoc*. However, both were primarily found at Western Crater, indicating that the *Eurotiomycetes* species on Mt. Erebus are either not lichenized or use unusual photosynthetic partners.

For the prokaryotes, we found that Exposed sites had greater variability in microbial communities than Subglacial sites. This is likely due to the large differences in temperature, water content (a consequence of different fumarolic activity levels), and pH (also attributed to different fumarolic activity levels (7)) between different Exposed samples (**Fig 1A**). One explanation for the variation in pH at high temperature sites could be that sulfur levels in the fumarolic gases may vary from fumarole to fumarole. Additionally, we note that Exposed sites also inherently contain more variability than Subglacial sites because Subglacial samples were usually collected from small pockets of loose soil amongst rocky terrain, while Exposed samples come from areas that are almost exclusively loose soil.

Also of note is the significantly higher abundance of ASVs from the phylum *Firmicutes* at Exposed Hot Soils compared to Tramway Ridge and Subglacial samples; one ASV from this phylum was also found to be indicator species for this subcategory (**Table S6**). The reason behind this is unclear, as most of the families identified are spore-forming, thermotolerant acidophiles, which should mean they would also be abundant at Tramway Ridge.

In our fungal data set, we observed clear differences in the types of fungi observed at different Exposed sites; however, determining ecological implications of these patterns is challenging given the phylogenetic novelty of many of the fungi in our data set. We did observe that the fungal ASVs predominant at Western Crater were mostly from clade 1, a highly divergent group of *Chytridiomycota* that showed similarity to fungi found in biological soil crusts in nearby Victoria Land (55). *Chytridiomycota* are best known for parasitic infections and degrading complex organic matter (56); their presence at Western Crater, which has extremely low organic content, is intriguing.

Finally, for our non-fungal eukaryotic data set, we discovered that amoebae ASVs were found primarily in Exposed samples (**Table S6**), although the most abundant amoeba ASV in our data set was a *Vermamoeba* species that was found in Helo Cave at 5% relative abundance. This particular species is known to be widespread and thermotolerant (57). Amoebae have previously been found on Mt. Erebus at Exposed sites (58) and viable amoeba cysts have been found in Arctic permafrost before (59), including similar groups to those found in some of our samples (especially Harry’s Dream).

#### Distinguishing features of Tramway Ridge sites: novel Nitrososphaeria

Overall, our expanded dataset of microbial ecosystems on Mt. Erebus further confirms that Tramway Ridge is an outlier across all other prokaryotic communities on Mt. Erebus, as has also been previously reported (7) (**Fig 2**). Despite conflicting past results about the role of temperature in structuring the microbial community within Tramway Ridge (4,7), our larger data set allowed us to confirm that there is a temperature threshold (55 °C) above which samples tend to cluster together away from other Tramway Ridge samples, indicating a strong upper temperature barrier. These higher temperature samples are characterized by a higher prevalence of *Deinococcota* (especially *Meiothermus*), which could simply indicate temperature preference.

The prokaryotic communities characteristic of Tramway Ridge have been discussed previously (4,5,7), with our extended sample set confirming these previous findings. Of particular note is confirmation that Tramway Ridge has characteristically high abundances of the archaeal phylum *Crenarchaeota*. One ASV (ASV_1) in particular was at high abundance, matching previous results (4,5). It was classified to the order SCGC AB-179-E04 (now belonging to the phylum *Thaumarchaeota* (60) and also known as *Thermoproteota* (61)) within the class *Nitrososphaeria*. This class is common in marine habitats (62), with members known for being autotrophic ammonia oxidizers (AOA); some deep-branching, thermophilic members lack the genes needed for ammonia oxidation (63–65). In particular, the order SCGC AB-179-E04, although under-studied, was recently found in high abundances (up to 79% of the prokaryotic community) in CO_2_ and H_2_-rich fumarolic vents from Asal-Ghoubbet rift, Republic of Djibouti (66). This is striking because CO_2_ and H_2_ have both been reported to be produced in high concentrations by Mt. Erebus (11,67). However, at the Asal-Ghoubbet site, no consumption of CO_2_ or H_2_ in aerobic or anaerobic incubations of the soils was observed (66). Even though we observed an enrichment in predicted aerobic ammonia oxidation at Exposed sites, (**Fig 7**), ASV_1 was not one of the 40 ASVs predicted to be involved in this functional category (all predicted ammonia oxidisers were from the classes *Nitrososphaera* or *Nitrocosmicus* of phylum *Crenarchaeota*). This indicates uncertainty in the functional prediction program based on the phylogenetic novelty of ASV_1. Clearly, further metagenomic and culturing studies are needed to investigate the function and phylogenetics of this highly abundant, novel archaeal species.

In our fungal data set, we note the identification of two fungal ASVs from the family *Herpotrichiellaceae* as indicator species for Tramway Ridge. Species from this family are generally rock-dwelling fungi, many of which are known to be opportunistic human pathogens (68).

### Hypothesis 2: Large numbers of unexpected fungi and taxonomically novel prokaryotes

Past studies have hypothesized that the biota of Mt. Erebus is enriched in taxonomically novel and potentially endemic organisms due to the unique geothermal features, geochemistry, and isolation of these sites. Three pieces of evidence point towards large proportions of taxonomically novel and potentially endemic biota on Erebus. First, a large proportion of fungal sequences in our data set had either no significant similarity (20 clades) or a very limited numbers of matches (three clades) to any DNA sequence in any database. It is likely that these sequences are highly novel groups of fungi, given the clear patterns we observed in abundance profiles of these novel clades at different sites (**S6 Fig**), the paucity of thermophilic fungi at low-organic sites in previous studies (69,70), and our finding that the novelty of clades increased at Exposed sites, where one would expect larger accumulations of dispersed spores and thus, less novel fungi. Indeed, a recent study of fungal diversity in biological soil crusts in nearby Victoria Land, Antarctica, also found a large proportion of putatively unknown fungal species (55). Second, we found a comparatively high (5.3 %) proportion of the prokaryotic community was Unassigned at the phylum level. This number can be dependent on the database used and when the database was accessed. Nonetheless, in general in soils, the number of Unassigned sequences at the phylum level is generally <1 % (71), while in hot springs, this number can be around 3 % (72). Finally, only a small proportion of the prokaryotic community across all site categories (25 % of ASVs, comprising 21.8 % of reads) had a predicted functional profile using the FAPROTAX dataset, due to the large number of ASVs that were unassignable at high taxonomic levels in our dataset, a number on the low end of previously published results (73,74). Taxonomic novelty is generally considered a marker for endemicity, but this hypothesis remains to be fully tested.

The detection in our data set of fungal species at high temperature sites is surprising since, although fungi are known to be capable of survival in many different types of extreme environment (e.g., extreme cold, dry, radiation, etc.), truly thermophilic fungi are rare (70). Thermophilic fungi are generally thought to be mesophiles that broadened their temperature range to take advantage of high nutrient situations (e.g., compost piles, bird nests, etc.) (69). However, we note that our results are only indicative of presence, not activity. This is important given that a previous study comparing the taxonomy of active vs. present fungi at Antarctic sites found a stark difference between the two communities (75). Additionally, a study examining growth tolerances of fungi isolated from sites within Victoria Land, Antarctica, including geothermally heated soils from Mt. Melbourne, only found one fungal isolate that could tolerate temperatures above 40 °C (76). Thus, our results should be treated as preliminary.

These results highlight the need for further studies, such as metagenomics, sequencing of full-length ribosomal sequences (for fungi), growth/activity measurements to identify active members of the fungal community, and/or culture-based studies, to truly understand the taxonomy and functions of the novel fungi/prokaryotes present. Indeed, a recent study reported the successful cultivation of a member of the WPS-2 phylum (now *Ca.* Eremiobacterota) from Warren Cave on the Mt. Erebus summit that is a novel aerobic anoxygenic photoheterotrophic bacterium (21). This highlights the importance of culturing in understanding the functions of highly divergent microorganisms.

### Hypothesis 3: Potential human influence on sites appears to be minimal

Although a past study identified large amounts of potentially human-associated fungal contaminants in Subglacial Erebus samples, the results from our data set on human impacts are more mixed. In fact, the presence of *Malasezzia* in some of our samples and in previously published Warren Cave samples does not necessarily stem from human activity on site, given their discovery in a broad range of habitats around the world (24) and their propensity to contaminate PCR reagents (25). We were particularly interested to find lignolytic fungi in our data set, given the almost complete lack of lignin-containing plants in Antarctica (77,78). The high identity matches suggest these species could have been introduced with construction material used to build the current or past huts on the Erebus summit or that these taxa employ mixotrophic strategies that have yet to be described. We also note with concern the presence of 192 ASVs from *Animalia* across all sites, although half of these have less than 10 reads in any sample. This highlights the need for a follow-up study using qPCR with specific primers and more stringent contamination protocols during sampling and sample processing, to eliminate the possibility of human contamination of samples during processing. We also note that the absence of other signatures of human contamination in our data set (e.g., human-associated bacteria) does not mean that those signatures are not present.

## Conclusion

Our survey of the biota inhabiting Mt. Erebus has given insights into the broad diversity of organisms inhabiting the southernmost active geothermal site in the world. Ice-free features on Mt. Erebus ranging from ice caves to steaming hot soils all harbor an impressive array of taxa, from novel Nitrososphaeria, to a wealth of novel fungi, to a range of photosynthetic microorganisms, with distinct communities at Subglacial sites, Exposed sites, and Tramway Ridge in particular. We hope this study will serve as a springboard for further, targeted, and in-depth studies of these unique and potentially endemic organisms. Additionally, we hope the community will continue to consider how to preserve these precious sites, as well as adhering to the rules that are already in place around access.

## Materials and Methods

### Sample site descriptions and soil collection

We collected 48 soil samples from 30 different sites on and near the summit of Mt. Erebus (3794 m elevation). These areas were generally divided into “Exposed” or “Subglacial” types. An overview of sample sites is seen in **Fig 1** and **S1 Fig** and details given in **Table S1**.

Exposed hot soils on Mt. Erebus are ice-free areas where the soil is thermally heated, to the extent that snow and ice do not accumulate. There are four main sites where Exposed hot soils are found: Tramway Ridge, Western Crater, Side Crater, and around the Main Crater. Tramway Ridge, protected under Antarctic Specially Protected Area (ASPA) 175 since 2002, lies at an altitude of 3350 m and is approximately 1.5 km northwest from the Main Crater. Tramway Ridge has numerous, actively steaming fumaroles, with the highest measured surface temperatures on Mt. Erebus (65 °C) (4,14). Steep temperature and chemical gradients occur around these fumarole openings due to the perennially cold air temperatures (less than −20 °C), with extensive moss beds and cyanobacterial mats (4,12,14). Approximately 1 km west from the Main crater lies Western Crater, a shallow depression that is highly exposed to maritime winds. Unlike Tramway Ridge, there are few actively steaming fumaroles at Western Crater and no visible moss or cyanobacterial mats, with soil temperatures only reaching 50 °C, and most hot soil spots are at least partially covered by ice hummocks (small transient snow enclosures) (see S1 Fig K for an example) (7). Side Crater, located immediately adjacent to Main Crater, is (currently) volcanically inactive; the bottom is covered with ice, but several areas of hot soil and actively steaming fumaroles are present, primarily on the steep eastern side of the crater (79). The active Main Crater, located at the summit of Mt. Erebus, is 250 m wide and 120 m deep, and harbors a persistent lava lake that produces regular strombolian eruptions (1,3). The walls of the Main Crater are inaccessible to sampling, but numerous hot soil sites exist around the rim of the Main Crater with high soil temperatures (>60 °C). In addition to these primary sites, small patches of Exposed hot soil can be found at various places around the summit (see Fig 1).

Subglacial hot soils on Mt. Erebus are mainly found in ice caves distributed around the summit (8,9). These caves are formed when gas emissions from the crust melt the ice covering the surface; the water re-freezes, creating a roof of ice (9). Many of the ice caves are interconnected and thought to constitute a ‘cave system’ rather than individual features (80). Although the soils in ice caves are generally similar in chemical characteristics, the ice caves vary in size, temperature, and light penetration (8,80). Reported soil temperatures from ice caves are so far generally lower than at Exposed sites (8). Subglacial hot soils other than ice caves can be found scattered around the summit, usually in the form of small ice hummocks (often < 0.5 m in diameter). While ice caves are quite persistent, ice hummocks are ephemeral structures since the ice often collapses between seasons.

Soil samples were collected during the field seasons (southern summer) of 2006-2007, 2009-2010, 2010-2011, 2011-2012, and 2019-2020. Before collecting the samples, soil temperatures were measured 0.5 cm below the soil surface using a CheckTempt1 thermometer (Hanna Instruments, USA). Prior to collection of soil, the top mat-like layer (if present) was removed and either discarded or used as a separate sample (“Mat” samples). Then, the topsoil (0-2 cm or 0-4 cm) below the mat was transferred into sterile centrifuge tubes using sterile spatulas; samples that are exceptions list the sample depth in the sample name (e.g., 1-4cm). All collected samples were kept frozen once collected and at −80 °C in the laboratory prior to processing.

### Geochemistry

Elemental composition of each soil sample was measured using Inductively Coupled Plasma–Mass Spectrometry (ICP-MS). Dried samples were ground using a Mixer Mill MM 400 (Retch, Haan, Germany) at a frequency of 27.5 1/s for 1 min and 50 sec. 1 g of each sample was weighed into a 50mL falcon tube, and acid digested using 10 mLs of a 1:6 dilution of HCl and 4 mLs of 1:3 dilution HNO3 with reflux at 80 °C for 30 minutes, or until no bubbles were observed upon gentle agitation. The digested contents was rinsed into a volumetric flask and diluted with milli q water to a final volume of 100 mL and left overnight. To remove soil particles, and reduce residual chloride concentration, samples were filtered using 0.45 μM membrane filter into 50mL volumetric flasks and made upto 50mL with Miliq water. 10mLs of each sample was then tipped into a 15mL falcon tube and 200 μL of pure nitric acid was added. Elemental composition was measured using an Agilent 8900 ICP-MS (Agilent Technologies, Santa Clara, California, USA) at the University of Waikato ICP-MS facility. The following elements were analyzed: Na, Mg, K, Ca, Sr, Ba, P, S, Se, Al, Tl, Pb, B, As, V, Cr, Mn, Fe, Co, Ni, Cu, Zn, Cd, and U. The results from ICP-MS were obtained from different instrument calibrations causing some variation in the isotopes of the elements measured and upper detection limits. Since Al, K, and Na reached their detection limits (5,000 ppm, 10,000 ppm, and 10,000 ppm, respectively) in some runs but not others, these elements were removed from some of the statistical analyses. Experimental and calibration controls were included in every run to allow for the identification of background elemental composition.

pH and conductivity were measured from a suspension of soil and milliQ water with a 1:2.5 w/w ratio. pH was measured using a HI2213 Basic pH/ORP/°C Meter/3-point calibration (Hanna Instruments, Rhode Island, USA) and adjusted for temperature. Conductivity was measured using a Thermo Scientific Orion 4-Star Benchtop pH/Conductivity Meter (Thermo Fisher Scientific, Massachusetts, USA).

Total moisture content in soil samples was estimated gravimetrically by drying 3 g of each soil sample in a drying oven at 105 °C. When the weight of a sample was unchanged over a 24 h period, the initial and final weight difference was converted into the moisture content percentage. The resulting dried samples were then prepared for total carbon and nitrogen measurements via grinding, as for ICP-MS analysis. Total carbon and nitrogen were measured from 1 g of each sample on a Vario EL Cube (Elementar, Langeselbold, Germany).

### DNA Extraction, 16S rRNA gene PCR, Sequencing, and Quality Control

A modification of the CTAB protocol developed previously (5) was used to perform triplicate DNA extractions on 0.7 – 0.9 g of soil. Additional extractions were performed for samples with low DNA yields. DNA from replicate extractions was pooled prior to analysis. The quality and concentration of the extracted DNA was checked using a Nanodrop DN-1000 spectrophotometer (Nanodrop Inc., Delaware, USA) and a Qubit 2.0 Fluorometer (Thermo Fisher Scientific).

16S rRNA amplicons were generated by triplicate 25 μL PCR reactions, using Ion Torrent Earth Microbiome fusion primers: 515YF (5’CCATCTCATCCCTGCGTGTCTCCGACTCAGXXXXXXXXXXXXXGATGTGYCAGC MGCCGCGGTAA3’) and 926R (5’CCACTACGCCTCCGCTTTCCTCTCTATGGGCAGTCGGTGATCCGYCAATTYMTT TRAGTTT3’). Each sample was amplified with a specific forward primer containing a unique barcode. The master mix contained 6 mM MgCl2, 0.24 mM dNTPs, 1.2× PCR buffer, 1.6×10-5 mg/μL BSA, 0.024 U/μL Taq polymerase (Thermo Fisher Scientific), 0.2 mM of each primer, and 1-3ng of sample DNA. PCR reactions were run on an Applied Biosystems ProFlex PCR System (Thermo Fisher Scientific) and the cycling conditions were: 94℃ for 3 min followed by 30 cycles of 94℃ for 45 sec, 50℃ for 1 min, and 72℃ for 1.5 min with a final extension of 72℃ for 10 min.

Triplicate PCR reactions were pooled, and 5 μL of each was visualized on 1% agarose gel. The Invitrogen SequalPrep Normalization kit (Thermo Fisher Scientific) was used to purify and normalize 25 μL of each amplicon, 2 μL of each normalized amplicon was pooled to make up the sequence library. The library was quantified using Qubit 2.0 Fluorometer (Thermofisher), and treated with Ion PGMTM Template IA 500 Kit (Thermo Fisher Scientific) before preparation for sequencing with using Ion PGMTM Hi-QTM View Sequencing Kit (Thermo Fisher Scientific). The library was added to an Ion 318TM Chip Kit v2 BC (Thermo Fisher Scientific) and sequenced on an Ion Torrent PGM (Thermo Fisher Scientific).

Resulting 16S rRNA gene sequences were filtered for quality and trimmed to generate amplicon sequence variants (ASVs) using the DADA2 v3.14 pipeline with the modifications recommended for Ion Torrent data (e.g., HOMOPOLYMER_GAP_PENALTY = –1, BAND_SIZE = 32) (81). The ASVs were assigned taxonomy using the SILVA database nr. 99 v138 (82). Sequences assigned to Eukaryotes, mitochondria, and chloroplast were removed. The R-packages Decipher (83) and phangorn (84) were used to generate an unrooted phylogenetic tree by the neighbor-joining method. See **Table S8** for number of reads at all stages in processing.

Samples taken from the different site categories had similar mean read counts (**S2 FigA-B**). However, two samples stood out; WC–10 and Crater 1 Mat had notably lower read counts (2,462 reads) and higher read counts (105,524), respectively, compared to other samples. Leaving these samples out from the dataset did not affect the calculated distributions in a PCoA analysis (**S2 FigC-D**). Lastly, rarefaction curves with the species richness were plotted (**S2 FigE**). The rarefaction curves all reached plateau, including WC-10 (**S2 FigF**). This indicates that the majority of the richness present in the samples has been covered, despite the low number of reads in the WC-10 sample.

### 18S rRNA gene and ITS region PCR, Sequencing, and Quality Control

Select samples (17) were chosen for ITS region (used for fungal classification) and 18S rRNA gene (used for non-fungal eukaryote classification) sequencing based on the type of geothermal feature to ensure representation from a variety of environments. DNA from these samples (also used for 16S rRNA gene sequencing) was sent to Omega Biosciences (Norcross, GA, USA) for amplification using the Earth Microbiome Project 18S rRNA gene or ITS primers without blocking sequences (85). They then underwent paired-end sequencing on an Illumina MiSeq v3 with 300 bp reads.

The resulting ITS and 18S rRNA gene sequences were adaptor trimmed using cutadapt (86) and quality checked using FastQC (87). Resulting trimmed sequences were processed using the DADA2 pipeline with default parameters for the respective workflows, except in the filterAndTrim step for both workflows, the following change was made: maxEE = c(3,5). For ITS sequences, taxonomy was assigned using the UNITE database v9 with singletons set as RefS (88). For 18S rRNA gene sequences, taxonomy was assigned using the SILVA v132 18S database formatted for DADA2 (89). For the 18S data set, all ASVs assigned to fungal groups were removed, while all ASVs assigned to non-fungal groups were removed from the ITS data set (see **Table S8** for number of reads at all stages in processing).

After generating the ITS ASVs with DADA2, we found that a large number of fungal sequences were poorly classified. We therefore conducted a phylogenetic analysis to identify putative clades of related ASVs and their taxonomic identities. We compiled a database consisting of (1) All ASV sequences, (2) The highest similarity sequence in the UNITE-SH database (88)) where that match had a percent identity > 90% over > 75% of the length, (3) Where there was no match in the UNITE-SH database, the highest similarity sequence in the UNITE+INSD database where that match had a percent identity > 90% over > 75% of the length, and (4) additional sequences from the NCBI nucleotide database in an iterative adaptive heuristic search strategy, attempting to improve the phylogenetic analysis and resolve identifications wherever possible. ASV and added sequences were aligned using default options in MAFFT (90); only references that were ITS1-5.8S-ITS2 were used to improve placement. Sequences were not trimmed. A nucleotide substitution model was determined using standard options in Jmodeltest (invgamma option) (91) and a phylogeny was then constructed using MrBayes (92) using default options.

From the phylogeny, high-level phylogenetic clades were identified based on >50% support. We named each clade on the basis of the lowest-numbered ASV present within that clade. As a measure of phylogenetic novelty, we also recorded the number of matches in the database where there were fewer than 5 at > 90% identity and > 75% of the ASV sequence length, and at > 80% identity and > 75% of the ASV sequence length. Lastly, for those clades or subclades where no ASVs matched sequences in UNITE databases, we searched the representative ASV sequences against the full NCBI nucleotide database (https://blast.ncbi.nlm.nih.gov/). Using the full NCBI database allowed inclusion of unidentified environmental sequences in the analysis, and also served as a check for any non-fungal sequences remaining in the data. Final taxonomic classification of ASVs in clades was based on the taxonomic classifications of reference sequences that fit into these phylogenetic clades.

### Statistical Analyses

Data analysis was conducted using R v. 4.2.2 and all plots were visualized using ggplot2 v. 3.4.0 (93) with beautification of figures in Inkscape (https://www.inkscape.org/).

A principal component analysis (PcoA) was performed to understand potential groupings of sites by first scaling the variables using the “scale” base function. Next, Euclidian distances were calculated using the “vegdist” function in the R package vegan v. 2.6.4 (94) and the PcoA calculated using the base “cmdscale” function. For testing physicochemical factors for significant differences between sites, outliers were first removed using the interquartile range method (95). Next, physicochemical parameters that were returned as significantly different between Subglacial and Exposed categories based on a two-sided t-test (*p*-values adjusted via Benjamini-Hochberg) were selected for further analysis (number of tests: 27). For this smaller list of parameters (n=12), significant differences between Site Subcategories were detected using a Krustal-Wallis test followed by Dunn’s test within ggstatsplot v. 0.12.3 (96), using the command ggbetweenstats (options: type = “nonparametric”, pairwise.comparisons = TRUE, p.adjust.method = “BH”) and *p*-value adjustment using a Benjamini-Hochberg correction (number of tests performed: 12).

Alpha diversity was measured with phyloseq v.1.40.0 using the Shannon index (97). Unconstrained beta diversity of the microbial community structures was investigated by center log-ratio transforming sequence abundances on a Unifrac distance matrix using microbiome v. 1.18.0 (98) and phyloseq, respectively. Different ways of categorizing sample sites were tested for statistical differences in the microbial community composition by using vegan to calculate a permutational analysis of variance (PERMANOVA) followed by an analysis of multivariate homogeneity (PERDISP) to verify that the dispersion of the tested samples did not differ significantly. The differential abundance test (trans_diff) within the microeco package was used to find specific taxa (at the phylum level) whose abundances differed significantly between either Site category or subcategory, using the random forest method (method = “rf”) and Benjamini-Hochberg correction. We used the R package indicspecies v. 1.7.14 to calculate Indicator Values for ASVs, with the command multipatt (99); only ASVs with strong indicator values (above 0.85) were included.

Environmental factors correlating with microbial community composition were identified using mantel tests within microeco, using spearman’s correlation and bray’s distance matrix and Benjamini-Hochberg corrected *p*-values: cal_ordination(method = “dbRDA”, use_measure = “bray”) cal_mantel(method = “spearman”, use_measure = “bray”, p_adjust_method = “BH”) The environmental factors that correlated significantly (*p* < 0.05, Benjamini-Hochberg correction) with microbial community structure were plotted using a distance-based Redundancy Analysis (dbRDA) within microeco (plot_ordination). Autocorrelations between environmental measures were investigated using the function “cor.test” from the package stats with spearman’s test; resulting p-values were adjusted with a Benjamini-Hochberg correction. To identify correlations between specific taxa and environmental factors, we used an RDA ($cal_ordination, method = “RDA”) at the phylum level within microeco, plotted using $plot_ordination. Correlations between taxa and environmental factors were calculated within microeco using a spearman’s correlation within the $cal_cor function with a Benjamini-Hochberg correction across all data (p_adjust_type = “All). The resulting heatmap was plotted using $plot_cor within microeco.

To predict functional profiles for the prokaryotic ASVs, we used a functional prediction program (FAPROTAX) (100) that uses taxonomic closeness to cultured strains with known functional traits in microeco. Functional predictions were calculated using the trans_func$new command, followed by pro.func$cal_spe_func(prok_database = c(“FAPROTAX”)). Correlations between predicted functional profiles and environmental factors were calculated with the $cal_cor using spearman’s test with a Benjamini-Hochberg correction for multiple tests across all data (p_adjust_type = “All”) on the resulting *p*-values.

To examine co-occurrence patterns between prokaryotes and eukaryotes, we constructed a co-occurrence network. First, to reduce compositional bias, we normalized each separate dataset (16S, 18S, and ITS) using center-log-ratio method (CLR) within the microbiome package v. 1.18.0. The data sets were then combined, with samples retained that had all three data sets represented. ASVs were then further trimmed to only retain ASVs that had at least a CLR of 4.5 in at least 3 samples, resulting in 228 ASVs (**Table S9**). Then, using NetCoMi v. 1.1.0 (101), a co-occurrence network was constructed, using the netConstruct command with Pearson as the calculation method, and “t-test” as the sparsification method with multiple testing adjustment using the adaptive Benjamini-Hochberg method: netConstruct(phy_erebus_all2, measure = “pearson”, sparsMethod = “t-test”, adjust = “adaptBH”, verbose = 3, seed = 63)

Clustering was performed using the fast and greedy option under the command netAnalyze, and singletons were removed prior to plotting.

## Acknowledgments

We especially recognize the logistical support from Antarctica New Zealand for fieldwork on Mt. Erebus and support in the field from Jon Tyler. We thank Drs. Alexis Marshall and Mafalda S. Baptista for support with quality control of sequencing data and scripts for ASV generation. This work was supported by the Royal Society of New Zealand (Marsden Grant 18-UOW-028 to SC, MS, IM, and CL).

## Data Availability

All R scripts used to analyze the data are available on Github at https://github.com/ThermophileResearchUnit/Erebus_survey_manuscript. All sequence data is available in DDBJ/EMBL/GenBank. 16S rRNA gene sequence data for almost all samples has been deposited under the accession KIGZ00000000. The version described in this paper is the first version, KIGZ01000000. 16S rRNA gene sequence data for samples TR1-*, TR2-*, and WC-* have been deposited under the accession KIGX00000000. The version described in this paper is the first version, KIGX01000000. 18S rRNA gene sequence data have been deposited under SUB14470052. ITS sequence data have been deposited under SUB14470177.

