## Supplementary Material for "Geothermal ecosystems on Mt. Erebus, Antarctica, support diverse and taxonomically novel biota"

**Supplementary Tables**

**Table S1.** Sample metadata and physicochemical data for all samples used in this study (separate table).

**Table S2.** Taxonomy and relative abundance information for all prokaryotic ASVs (separate table).

**Table S3.** Taxonomy and relative abundance information for all fungal ASVs (separate table).

**Table S4.** Taxonomy and relative abundance information for all non-fungal eukaryotic ASVs (separate table).

**Table S5.** Fungal clade identities (based on phylogenetic approach) and statistics on size, novelty, and probably taxonomy (highest possible taxonomic classification based on reference sequence grouping) of clades.

| Clade Number | # ASVs | #UNITE matches >90% ID | #UNITE matches >80% ID | #INSD matches >90% ID | # INSD matches >80% ID | # NCBI matches >90% ID | # NCBI matches >80% ID | Identity |
|---|---|---|---|---|---|---|---|---|
| **Clade_1** | 3 | 0 | 4 | 1 | 4 | NA | NA | Chytridiomycota |
| **Clade_2** | 1 | 1 | 2 | ≥5 | ≥5 | NA | NA | Ascomycota |
| **Clade_3** | 5 | 0 | 0 | 0 | 0 | 0 | 0 | Fungi |
| **Clade_4** | 1 | 0 | 0 | 1 | 1 | 1 | 1 | Fungi |
| **Clade_5** | 14 | ≥5 | ≥5 | ≥5 | ≥5 | NA | NA | Ascomycota |
| **Clade_6** | 1 | 0 | 0 | 0 | 0 | 0 | 0 | Pezizomycotina |
| **Clade_7** | 4 | ≥5 | ≥5 | ≥5 | ≥5 | NA | NA | Leotiomycetes |
| **Clade_9** | 2 | 0 | 0 | 1 | 1 | 5 | 5 | Ascomycota |
| **Clade_10** | 3 | 0 | ≥5 | ≥5 | ≥5 | NA | NA | Herpotrichiellaceae |
| **Clade_14** | 1 | ≥5 | ≥5 | ≥5 | ≥5 | NA | NA | Stereum |
| **Clade_15** | 4 | ≥5 | ≥5 | ≥5 | ≥5 | NA | NA | Herpotrichiellaceae |
| **Clade_16** | 2 | 0 | 0 | 0 | 0 | 0 | 0 | Eukaryota |
| **Clade_19** | 3 | 1 | 3 | ≥5 | ≥5 | 5 | 5 | Eukaryota |
| **Clade_20** | 7 | ≥5 | ≥5 | ≥5 | ≥5 | NA | NA | Filobasidiales |
| **Clade_23** | 9 | ≥5 | ≥5 | ≥5 | ≥5 | NA | NA | Dothideomycetes |
| **Clade_25** | 1 | 0 | 0 | 0 | 0 | 0 | 0 | Fungi |
| **Clade_26** | 5 | ≥5 | ≥5 | ≥5 | ≥5 | NA | NA | Agaricales |
| **Clade_28** | 3 | 0 | 0 | 0 | 1 | 0 | 1 | Fungi |
| **Clade_29** | 11 | ≥5 | ≥5 | ≥5 | ≥5 | NA | NA | Malasseziaceae |
| **Clade_30** | 3 | 3 | ≥5 | ≥5 | ≥5 | 5 | 5 | Hydnaceae |
| **Clade_31** | 12 | 0 | 0 | 2 | 3 | 1 | 1 | Fungi |
| **Clade_32** | 3 | ≥5 | ≥5 | ≥5 | ≥5 | NA | NA | Leotiomycetes |
| **Clade_34** | 6 | ≥5 | ≥5 | ≥5 | ≥5 | NA | NA | Pleosporales |
| **Clade_41** | 2 | 2 | ≥5 | ≥5 | ≥5 | 2 | 5 | Parmeliaceae |
| **Clade_42** | 6 | ≥5 | ≥5 | ≥5 | ≥5 | NA | NA | Ascomycota |

| | | | | | | | | |
|---|---|---|---|---|---|---|---|---|
| **Clade_44** | 3 | 3 | 4 | ≥5 | ≥5 | NA | NA | Malasseziaceae |
| **Clade_48** | 2 | 0 | 0 | 0 | 0 | 0 | 0 | Fungi |
| **Clade_49** | 1 | 0 | 0 | 0 | 0 | 0 | 0 | Chytridiomycota |
| **Clade_54** | 2 | ≥5 | ≥5 | ≥5 | ≥5 | NA | NA | Cystobasidiomycetes |
| **Clade_55** | 4 | ≥5 | ≥5 | ≥5 | ≥5 | NA | NA | Sporidiobolaceae |
| **Clade_56** | 1 | 0 | 0 | 0 | 0 | 0 | 0 | Eukaryota |
| **Clade_57** | 1 | 0 | 0 | 0 | 0 | 0 | 0 | Eukaryota |
| **Clade_63** | 6 | 3 | ≥5 | ≥5 | ≥5 | 1 | 0 | Polyporales |
| **Clade_69** | 2 | 0 | 0 | 0 | 0 | 0 | 0 | Ascomycota |
| **Clade_70** | 1 | 2 | 3 | ≥5 | ≥5 | NA | NA | Malasseziaceae |
| **Clade_73** | 1 | 0 | ≥5 | 0 | ≥5 | 2 | 5 | Fungi |
| **Clade_74** | 2 | 0 | 0 | 0 | 1 | 5 | 5 | Basidiomycota |
| **Clade_75** | 6 | ≥5 | ≥5 | ≥5 | ≥5 | NA | NA | Ascomycota |
| **Clade_83** | 2 | 0 | 0 | 0 | 0 | 1 | 1 | Fungi |
| **Clade_84** | 6 | 0 | 0 | 0 | 0 | 5 | 5 | Fungi |
| **Clade_85** | 1 | 0 | 0 | 0 | 0 | 0 | 1 | Rozellomycota |
| **Clade_88** | 1 | 0 | ≥5 | 0 | ≥5 | NA | NA | Ascomycota |
| **Clade_89** | 4 | 1 | ≥5 | 4 | ≥5 | NA | NA | Dothideomycetes |
| **Clade_91** | 1 | 2 | 4 | ≥5 | ≥5 | NA | NA | Fungi |
| **Clade_93** | 2 | 0 | 0 | 0 | 0 | 0 | 0 | Eukaryota |
| **Clade_100** | 1 | 2 | 3 | ≥5 | ≥5 | NA | NA | Sterigmatomyces |
| **Clade_107** | 1 | 1 | 2 | 2 | 3 | NA | NA | Herpotrichiellaceae |
| **Clade_108** | 2 | ≥5 | ≥5 | ≥5 | ≥5 | NA | NA | Pleosporales |
| **Clade_110** | 2 | ≥5 | ≥5 | ≥5 | ≥5 | NA | NA | Fungi |
| **Clade_111** | 1 | 1 | ≥5 | ≥5 | ≥5 | NA | NA | Rozellomycota |
| **Clade_112** | 3 | ≥5 | ≥5 | ≥5 | ≥5 | NA | NA | Dothideomycetes |
| **Clade_113** | 2 | 0 | 0 | 0 | 0 | 1 | 3 | Fungi |
| **Clade_121** | 1 | ≥5 | ≥5 | ≥5 | ≥5 | NA | NA | Ascomycota |
| **Clade_125** | 1 | 1 | 1 | ≥5 | ≥5 | NA | NA | Malasseziaceae |
| **Clade_127** | 1 | 0 | 0 | 0 | 0 | 5 | 5 | Fungi |
| **Clade_135** | 2 | ≥5 | ≥5 | ≥5 | ≥5 | NA | NA | Endogonomycetes |
| **Clade_136** | 1 | 1 | 1 | ≥5 | ≥5 | 2 | 4 | Rozellomycota |
| **Clade_137** | 1 | 4 | ≥5 | ≥5 | ≥5 | NA | NA | Ustilaginaceae |
| **Clade_141** | 2 | 0 | 0 | 0 | 1 | 2 | 5 | Fungi |
| **Clade_142** | 1 | ≥5 | ≥5 | ≥5 | ≥5 | NA | NA | Fungi |
| **Clade_144** | 1 | 1 | 1 | ≥5 | ≥5 | NA | NA | Schizoporaceae |
| **Clade_147** | 1 | 0 | 1 | 2 | 3 | NA | NA | Malasseziaceae |
| **Clade_156** | 2 | 2 | 3 | ≥5 | ≥5 | NA | NA | Exobasidiomycetes |
| **Clade_157** | 2 | 0 | 0 | 0 | 0 | 0 | 0 | Eukaryote |
| **Clade_161** | 1 | 0 | 0 | 0 | 0 | 0 | 0 | Ascomycota |
| **Clade_162** | 1 | 2 | 4 | 4 | ≥5 | NA | NA | Chytridiomycota |

| | | | | | | | | |
|---|---|---|---|---|---|---|---|---|
| **Clade_168** | 1 | 0 | 0 | 0 | 0 | 0 | 0 | Fungi |
| **Clade_174** | 1 | ≥5 | ≥5 | ≥5 | ≥5 | NA | NA | Botryosphaeriaceae |
| **Clade_181** | 1 | 0 | 0 | 0 | 0 | 0 | 0 | Eukaryote |
| **Clade_186** | 1 | 0 | 1 | 0 | 1 | 0 | 4 | Schizoporaceae |
| **Clade_187** | 1 | 0 | 0 | 0 | 0 | 0 | 0 | Eukaryote |
| **Clade_188** | 1 | 1 | 1 | 4 | 4 | 0 | 0 | Fungi |
| **Clade_189** | 1 | ≥5 | ≥5 | ≥5 | ≥5 | NA | NA | Glossaceae |
| **Clade_190** | 2 | 0 | 1 | 0 | 2 | 0 | 2 | Fungi |
| **Clade_192** | 1 | 0 | 0 | 0 | 0 | 0 | 0 | Eukaryote |
| **Clade_193** | 1 | 0 | 0 | 0 | 0 | 5 | 5 | Fungi |
| **Clade_194** | 1 | 1 | 3 | ≥5 | ≥5 | NA | NA | Lachnocladiaceae |
| **Clade_196** | 1 | 0 | 0 | 0 | 0 | 0 | 0 | Eukaryote |
| **Clade_197** | 1 | 1 | 1 | ≥5 | ≥5 | NA | NA | Chytridiomycota |
| **Clade_198** | 1 | 0 | 0 | 0 | 0 | 0 | 5 | Fungi |
| **Clade_200** | 1 | 0 | 0 | 0 | 0 | 0 | 0 | Eukaryote |
| **Clade_201** | 1 | 0 | 0 | 0 | 0 | 5 | 5 | Eukaryote |
| **Clade_205** | 1 | 4 | ≥5 | ≥5 | ≥5 | NA | NA | Fungi |
| **Clade_208** | 1 | 0 | 0 | 0 | 0 | 0 | 0 | Eukaryote |
| **Clade_209** | 1 | 0 | 0 | 0 | 0 | 0 | 0 | Fungi |
| **Clade_212** | 1 | 0 | 0 | 0 | 0 | 1 | 3 | Eukaryote |
| **Clade_214** | 1 | 1 | 4 | ≥5 | ≥5 | NA | NA | Herpotrichiellaceae |

**Table S6.** Results of indicator species analysis to identify species that are unique at the amplicon sequence variant (ASV) level for each Site Subcategory, either within Prokaryotes, Fungi, or non-fungal eukaryotes (separate table).

**Table S7.** Correlations between environmental factors across all sites. Only correlations with significant adjusted *p*-values and with correlation coefficients above 0.7 are shown. Also excluded are elements that were not identified as correlating significantly with the microbial community (see Table 1). All *p*-values were adjusted using a Benjamini-Hochberg correction. Number of tests performed: 351.

| **Variable 1** | **Variable 2** | **Correlation Coefficient** | ***p*-value** | **Adjusted *p*-value** |
|---|---|---|---|---|
| Temperature | Nitrogen | 0.748 | 8.53E-07 | 1.50E-05 |
| Conductivity | GWC | 0.764 | 3.68E-07 | 9.23E-06 |
| Carbon | Nitrogen | 0.925 | 4.86E-14 | 1.70E-11 |
| Carbon | GWC | 0.729 | 2.19E-06 | 3.15E-05 |
| Nitrogen | GWC | 0.774 | 2.00E-07 | 5.84E-06 |
| Mg | Ca | 0.875 | 1.82E-07 | 5.80E-06 |
| Ca | Sr | 0.806 | 6.99E-07 | 1.44E-05 |

**Table S8.** Tracking reads from samples through the DADA2 pipeline and post-processing steps across all domains.

**Table S9.** List of ASVs included in the co-occurrence network and corresponding abundance (center-log ratio adjusted) values (separate table).

**Supplementary Figures**

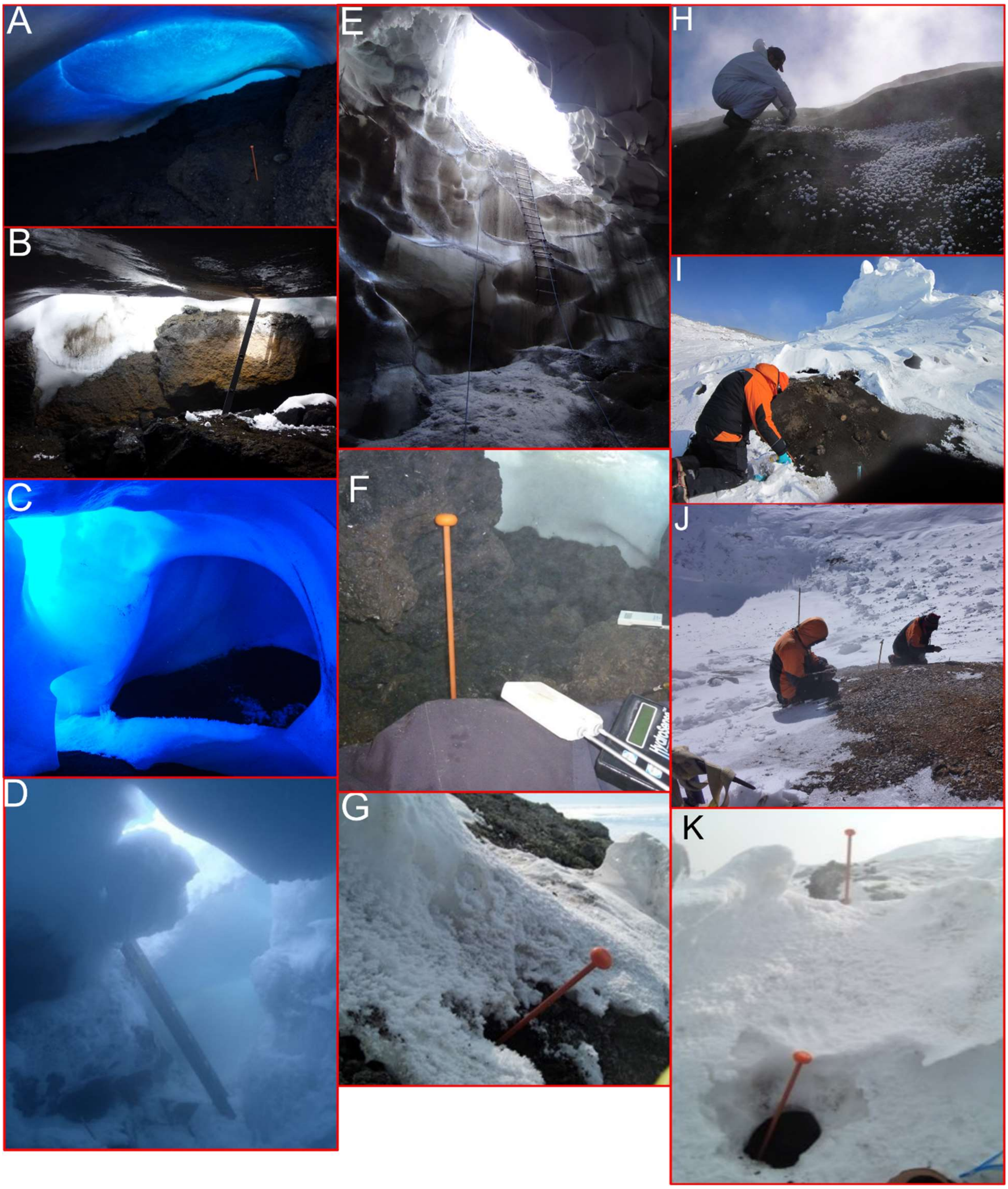


**Figure S1** Representative images of selected sampling sites. (A) Hut Cave interior. (B) Entrance to Helo Cave. (C) Entrance to 22 Blue Cave, as well as the sampling site (mostly dark). (D) Entrance to Harry’s Dream cave. (E) Entrance to Warren Cave. (F) Sampling site for Subglacial hot soil 8; the top of the ice hummock was removed to access the hot soil underneath for sampling.

(G) Sampling site for Exposed hot soil 6. (H) Sampling site at Tramway Ridge. (I) Sampling site at Western Crater. (J) One of the sampling sites in Side Crater; in the background is the floor of Side Crater. (K) Sampling site for Exposed hot soil 7 (foreground). Images are courtesy of Ian McDonald, Craig Cary, Craig Herbold, and Matthew Stott.

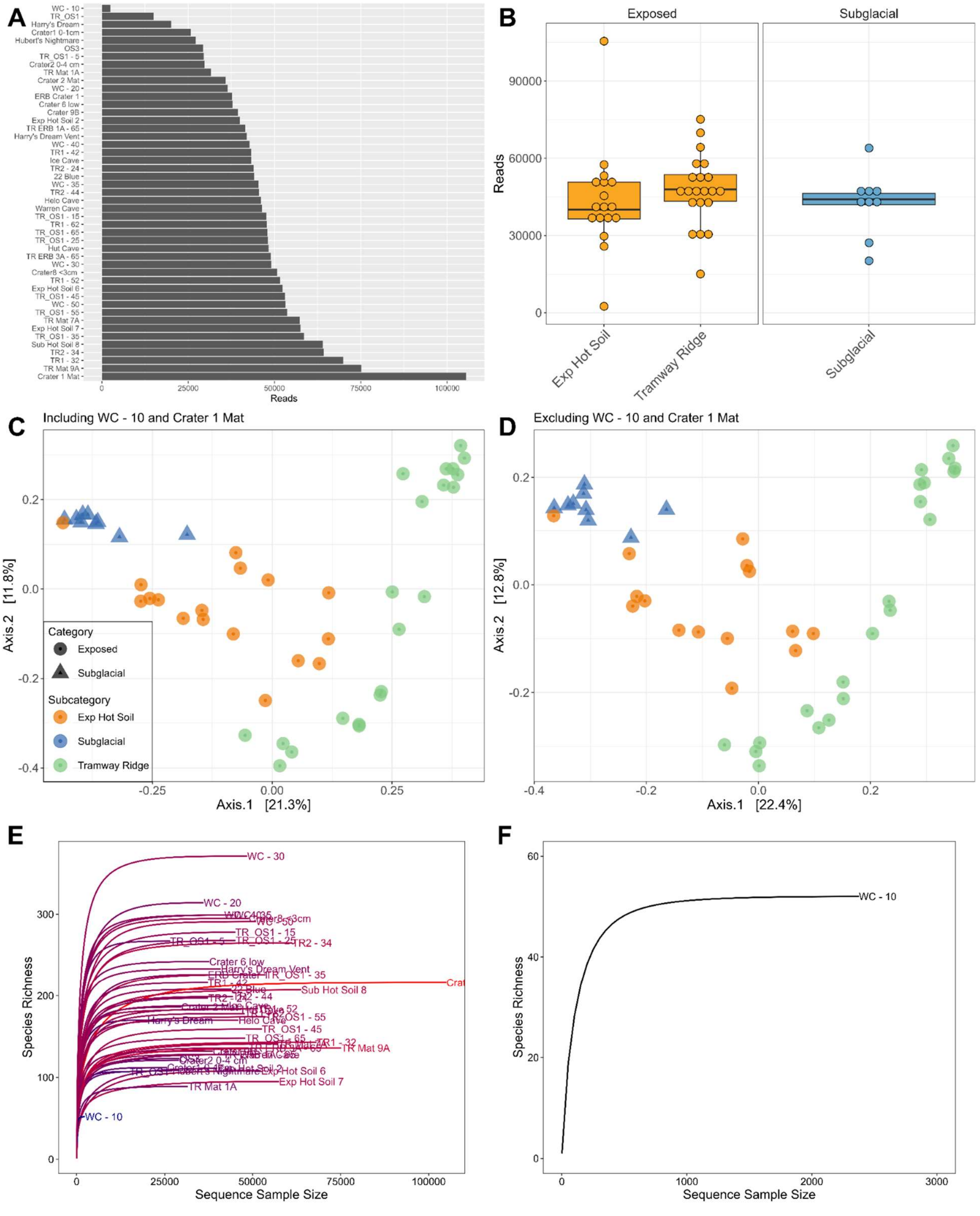


**Figure S2.** Initial assessment of amplicon sequencing data ensuring comparability between samples across sites of Mt Erebus. (A) The number of sequencing reads varied by two magnitudes across the different samples with 'Western Crater - 10' (WC - 10) having the least (2,462 reads) and 'Crater 1 Mat' having greatest number of reads (105,524). However, (B) the median number of reads between the different groups of samples did not differ significantly (pairwise wilcox test; $p > 0.5$). Furthermore, removing the Crater 1 Mat and Western Crater - 10 reads from beta diversity assessments only marginally impacted associations as depicted in the NMDS plots; (C) was generated with WC-10 & Crater 1 Mat, and (D) without. Lastly, plotting the species richness against the sequence sample size showed that the rarefraction curves for all samples (E), including Western Crater - 10 (F), reached plateau, meaning that the species richness was unlikely to be impacted substantially if more reads had been included.

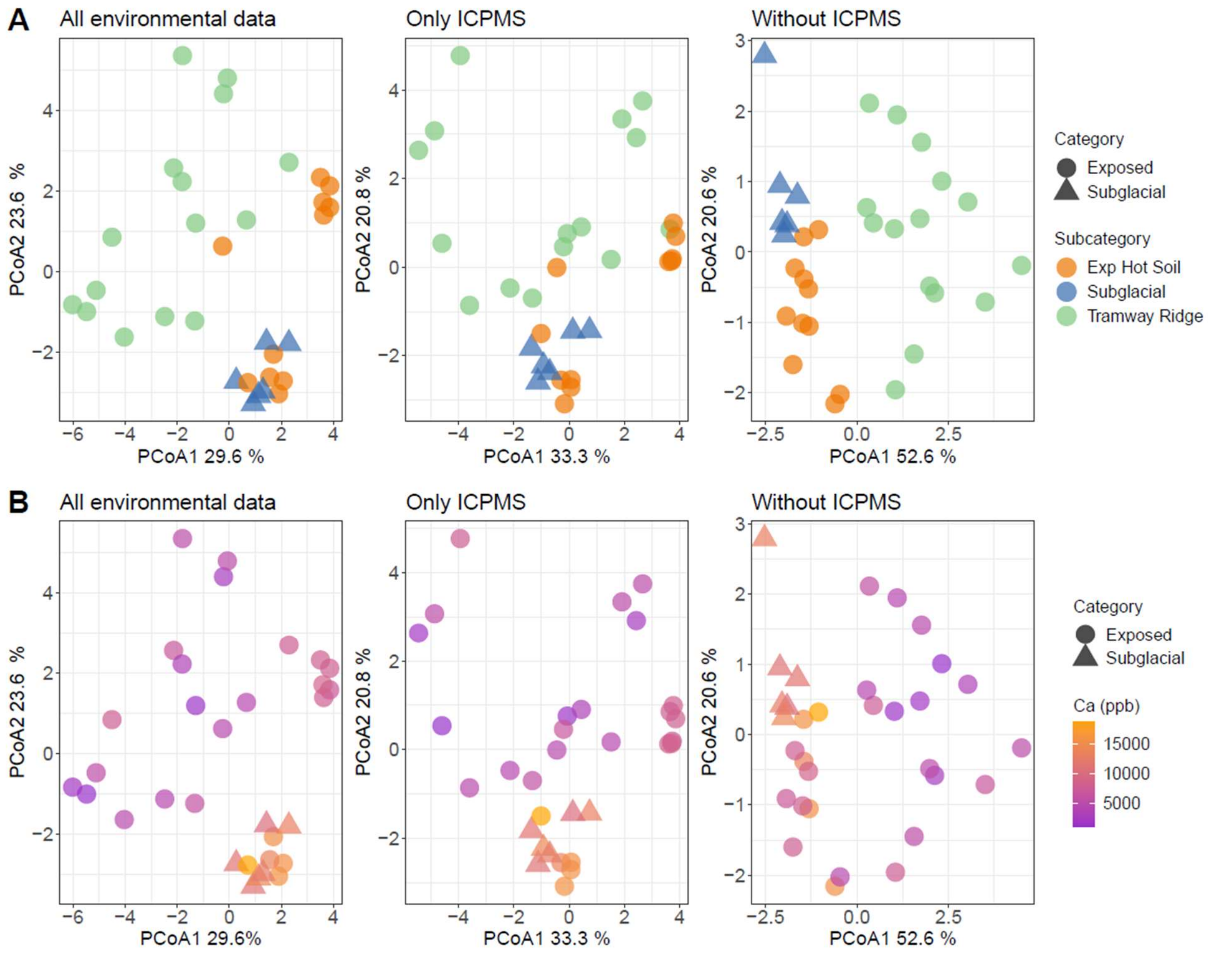


**Figure S3.** PCoA analyses of all physicochemical data, soil elemental composition, and physicochemical data minus elemental data from sample sites across Mt Erebus. The first column includes all the environmental data including pH, electroconductivity, temperature, gravitational water content, total carbon, total nitrogen, and ICP-MS elemental data. The second and third columns show only the elemental ICP-MS data and the physiochemical data excluding the elemental ICP-MS data. (Row A) samples are colored by its subcategory, i.e. Exposed Hot Soil (Exp Hot Soil), Subglacial, or a Tramway Ridge sample. (Row B) Samples are colored by calcium concentration (ppb).

**Figure S4**. Unrooted, Bayesian phylogenetic tree of fungal ASVs from our data set, as well as related reference sequences from NCBI, UNITE, and INSD databases (see Methods for full description). (Figure S4 is in a separate file).

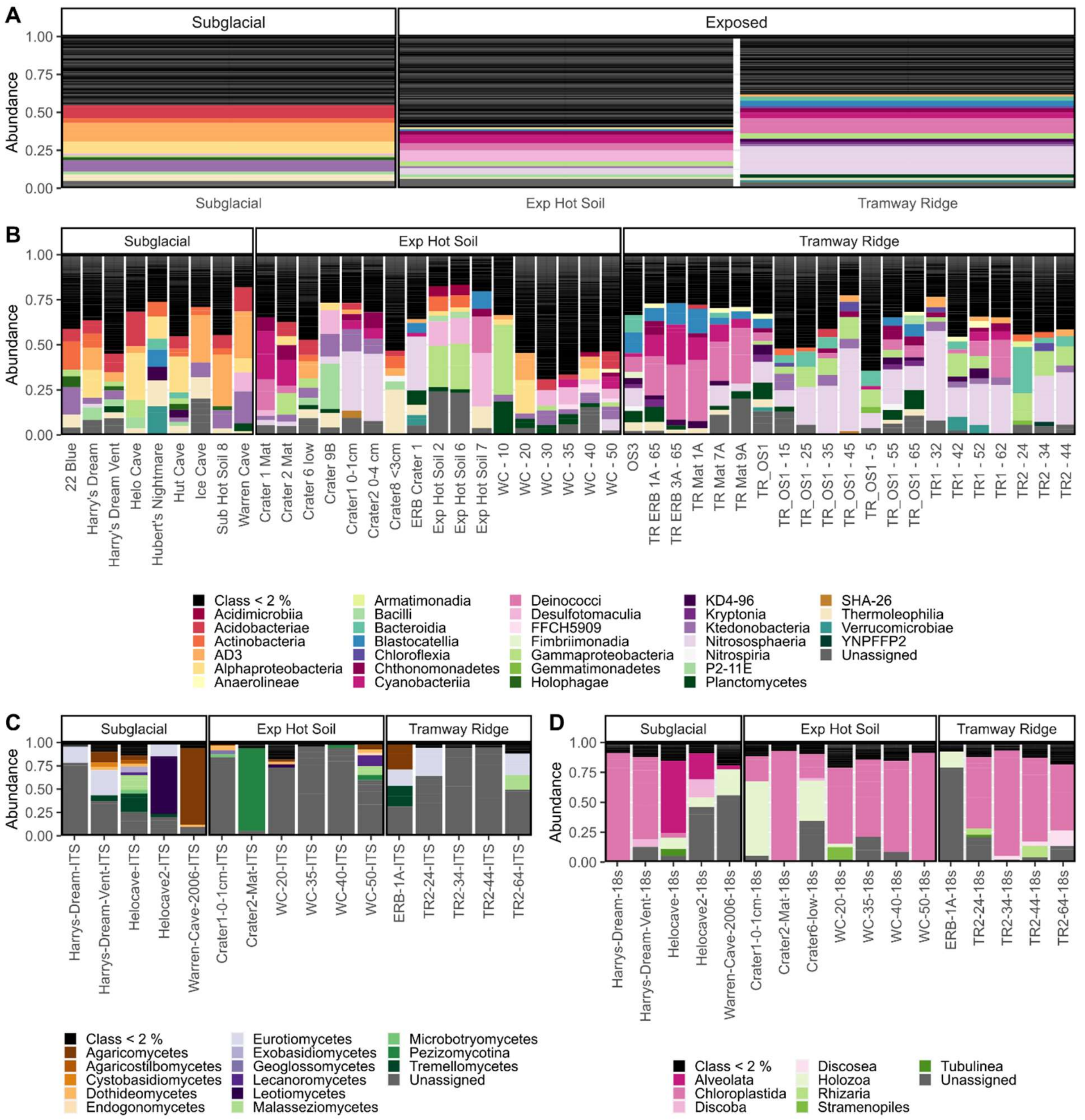


**Figure S5.** Barplots of taxonomic richness at the class level for (A) prokaryotes, (B) fungi, and (C) non-fungal eukaryotes from Subglacial, Exposed hot soil and Tramway Ridge sites on Mt Erebus.

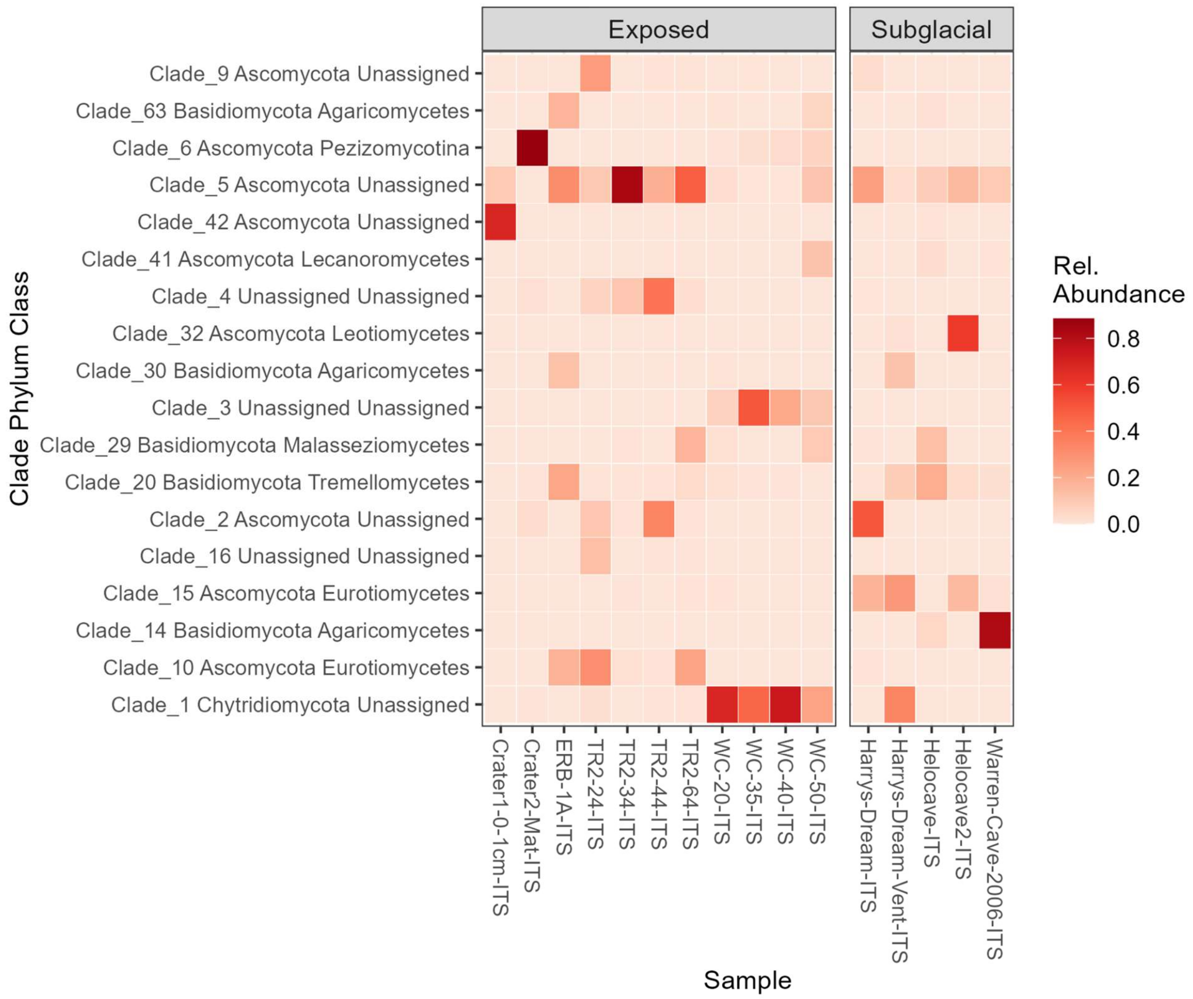


**Figure S6.** Abundance profiles of fungal clades (relative abundance > 10% in at least one sample) from Exposed and Subglacial sample sites on Mt Erebus. TR: Tramway Ridge. WC: Western Crater.

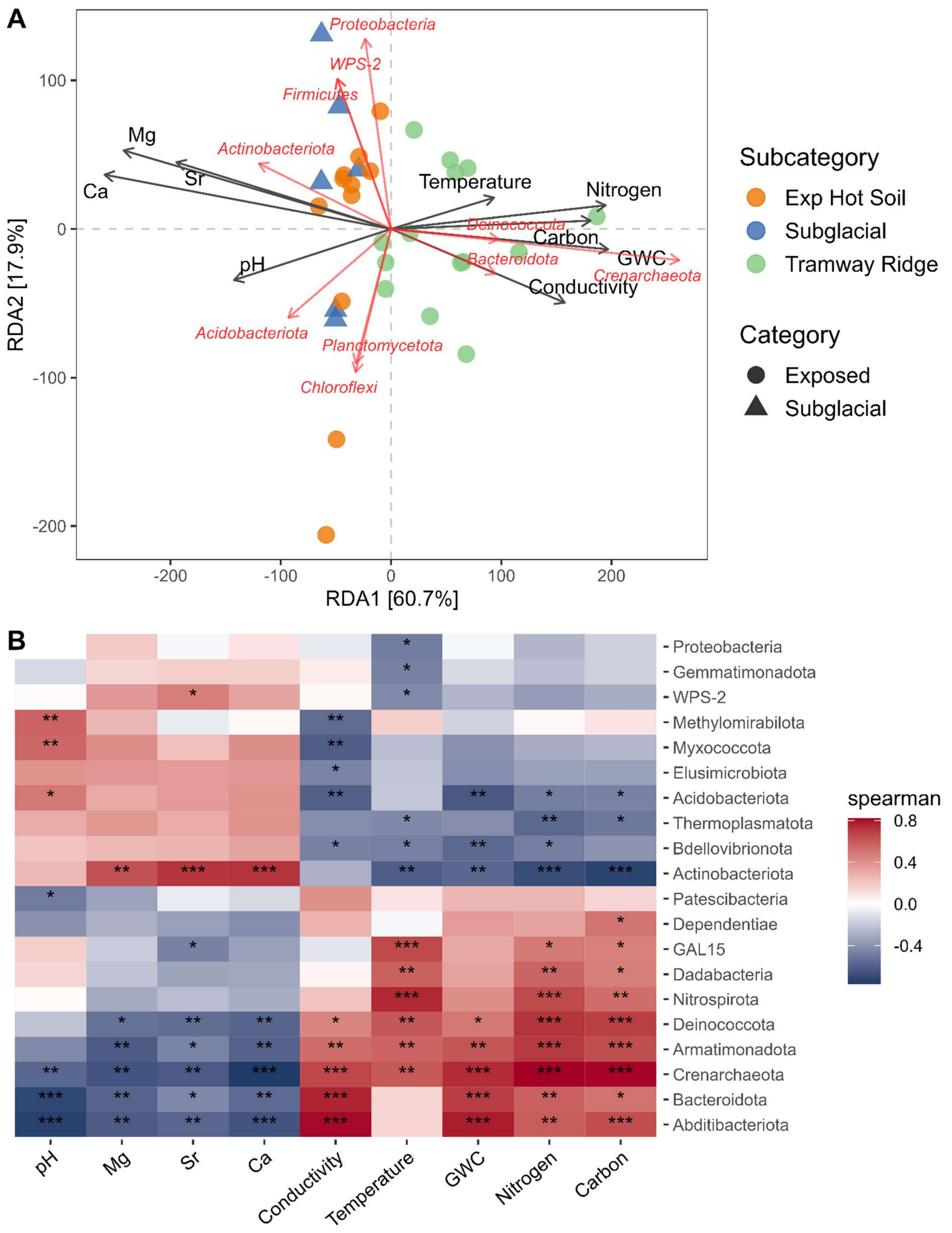
A
RDA2 [17.9%]
RDA1 [60.7%]
100
0
-100
-200
-200
-100
0
100
200
Proteobacteria
WPS-2
Firmicutes
Actinobacteriota
Mg
Sr
Ca
pH
Temperature
Nitrogen
Deinococcota
Carbon
GWC
Bacteroidota
Crenarchaeota
Conductivity
Acidobacteriota
Planctomycetota
Chloroflexi
Subcategory
Exp Hot Soil
Subglacial
Tramway Ridge
Category
Exposed
Subglacial
B
Proteobacteria
Gemmatimonadota
WPS-2
Methylomirabilota
Myxococcota
Elusimicrobiota
Acidobacteriota
Thermoplasmatota
Bdellovibrionota
Actinobacteriota
Patescibacteria
Dependentiae
GAL15
Dadabacteria
Nitrospirota
Deinococcota
Armatimonadota
Crenarchaeota
Bacteroidota
Abditibacteriota
pH
Mg
Sr
Ca
Conductivity
Temperature
GWC
Nitrogen
Carbon
spearman
0.8
0.4
0.0
-0.4

**Figure S7.** (A) Redundancy analysis of prokaryotic communities from Tramway Ridge, and Exposed and Subglacial soils on Mt Erebus with physicochemical factors and taxa at the phylum level incorporated as explanatory variables. (B) Spearman Rank correlation heatmap of physicochemical parameters and prokaryotic phyla. GWC: gravitational water content (moisture). The significance stars correspond to the following adjusted *p*-values (Benjamini-Hochberg correction): * = 0.05, ** = 0.01, *** = 0.001, **** = 0.0001. Number of tests performed: 261.
