## Supplementary figures and images for "Geothermal ecosystems on Mt. Erebus, Antarctica, support diverse and taxonomically novel biota"

### Figure S4

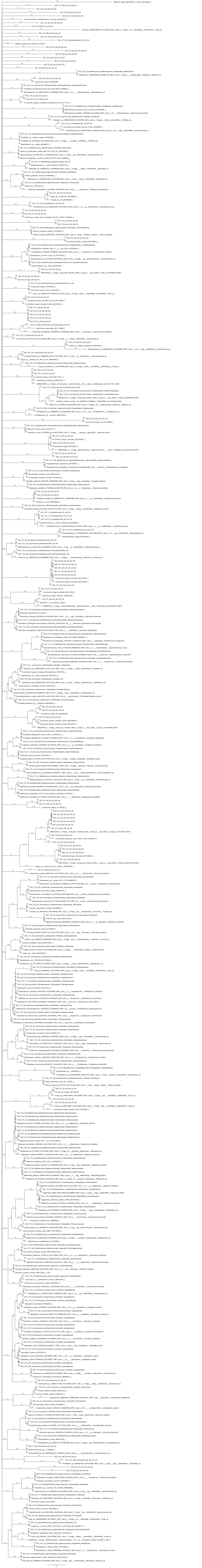
